# Linking intercontinental biogeographic events to decipher how European vineyards escaped Pierce’s disease

**DOI:** 10.1101/2024.05.04.592514

**Authors:** Eduardo Moralejo, Àlex Giménez-Romero, Manuel A. Matías

## Abstract

Unlike most grapevine diseases of American origin, the vector-borne bacterium *Xylella fastidiosa* (Xf) responsible for Pierce’s disease (PD) has not yet spread to continental Europe. The reasons for this lack of invasiveness remain unclear. Here, we present phylogenetic, epidemiological and historical evidence to explain how European vineyards escaped Xf. Using Bayesian temporal reconstruction, we show that the export of American grapevines to France as rootstocks to combat phylloxera (∼1872-1895) preceded the spread of the Xf grapevine lineage in the US. In the dated tree, the time of the most recent common ancestor places the introduction of Xf into California around 1875, which agrees with the emergence of the main PD outbreak and the onset of its expansion into the southeastern US around 1895. We also show that between 1870 and 1990, climatic conditions in continental Europe were mostly below the threshold for PD epidemics. This lack of spatiotemporal concurrence between factors that could facilitate the establishment of the Xf grapevine lineage would explain the historical absence of PD in continental Europe. However, our model indicates that there has been an inadvertent expansion of risk in southern European vineyards since the 1990s, which is accelerating with global warming. Our temporal approach identifies the biogeographic conditions that have so far prevented PD, and gives continuity to predictions of increased risk in important southern European wine-producing areas under a forthcoming scenario of +2 and +3°C temperature increases.

## Introduction

No crop better illustrates the dangers associated with global plant trade than grapevine^1^. During the 19th century, the accidental introduction of powdery mildew (*Uncinula necator*) ∼ 1845^2^ into Europe from exported North American wild *Vitis* species and the subsequent search for resistant germplasm led to the arrival of other native pests and pathogens from the American continent— e.g. phylloxera (*Daktulosphaira vitifoliae*) ∼1866^3^ and downy mildew (*Plasmopara viticola*) ∼1878^4^. In less than two decades, the entire European wine industry was devastated by the phylloxera crisis, while downy and powdery mildew have since become endemic diseases causing major economic losses.

The global spread of American grapevine’s pathogens and pests was triggered by the lack of proper sanitary control of plant propagating material, coupled with improvements in shipping and rail transport in the 19th century^5^. Surprisingly, the bacterium *Xylella fastidiosa* (Xf), the causal agent of Pierce’s disease (PD) and a limiting factor for viticulture in the southeastern US, presumably was not exported to Europe. This absence is even more puzzling when one considers that Xf is a vascular pathogen that is normally transmitted by xylem sap-feeding insects, but can also be transmitted by grafting^6^, as are many viruses, viroids and phytoplasmas introduced by this route^7^. It was initially speculated that the absence of PD in Europe might be due to a lack of competent vectors^8^ and/or to non-conducive climate conditions^9^, but despite substantial advances in the understanding of Xf biology, no compelling explanation has yet been provided.

PD was first described in 1884 in the vineyards of Anaheim, California. In less than a decade, the disease had spread throughout southern California and leaped to Napa Valley in northern California^10^. In his study, Newton Pierce documented in detail the unprecedented outbreak that ruined the prosperous viticulture of southern California. Although a grapevine ‘degeneration disease’ similar to PD had been described in Florida since around 1895^11^, the first unequivocal identification of PD in the southeastern US waited until 1951^11^. Additionally, some aspects of the aetiology of the disease remained obscure for a long time; in 1942 it was demonstrated that the pathogen is transmitted by insects belonging to the sharpshooter leafhoppers (Hemiptera: Cicadellinae) and spittlebugs (Hemiptera:Cercopoidae)^12^, whereas the bacterial nature of the disease remained unclear until 1978^13^.

During the 20th century, the endemic southeastern US origin hypothesis for grapevine-associated Xf strains prevailed due to the greater resistance of native American grapevines compared to European *V. vinifera* cultivars^14^. However, early genetic analyses using multi-locus sequence typing (MLST) revealed very little genetic variation among Xf subspecies *fastidiosa* isolates from grapevines across the US, suggesting a recent introduction and spread as a clonal complex^15,16^ (hereafter referred to as Xf_PD_). Shortly before, Xf_PD_ strains were also demonstrated to cause almond leaf scorch in California and to infect other hosts^17^. To date, Xf_PD_ has been found in Mexico^18^, Costa

Rica^19^, Taiwan^20^, Majorca, Balearic Islands, Spain^21^ and Israel^22^. Recent phylogenetic analysis of the global population of Xf_PD_ on grapevine using whole genome sequences has shown that it consists of two major clades, referred here as the Western US Clonal Complex (WUCC) and the Eastern US Clonal Complex (EUCC). Both major clades show small genetic distances and are nested within the larger genetic diversity of Xf subsp. *fastidiosa* isolates from Central America, the most likely centre of origin of the subspecies *fastidiosa* ^23–25^.

PD has long been considered a potential threat to the European wine industry. Such concerns have increased following the first detection of Xf in Europe in 2013 as the causal agent of olive quick decline syndrome in Puglia, Italy^26^. Since then, several sequence types belonging to different Xf subspecies have been detected in Corsica and the Provence-Alpes-Côte d’Azur (PACA) region of France, the Balearic Islands, Alicante (Spain), Portugal and Tuscany^27^. All these outbreaks have in common that the insect *Philaenus spumarius* plays a key and almost unique role in the transmission of Xf-associated diseases in Europe, including PD^28,29^. Today, the three subspecies of Xf, *fastidiosa*, *multiplex* and *pauca*, occupy a small area of their potential geographical distribution in Europe according to recently published species distribution models (SDMs) ^30,31^. However, these predictions have been criticised for disregarding both epidemiological dynamics and vector distribution as limiting factors of invasiveness. A much more restricted potential distribution of PD has recently been proposed using mechanistic models that incorporate the relationship between climatic conditions, pathogen growth *in planta*, and vector distribution and abundance^32^. Such PD risk maps based on epidemiological models driven by climate data are more reliable and allow accurate projections under future climate change scenarios^33^.

Despite the broad outlines of the evolutionary history of Xf_PD_ are known^24^, a chronological estimate of its spread within the US has not yet been attempted. Additionally, little attention has been paid to the conflicting results between the origin of PD genetic resistance in native wild *Vitis* species and the biogeography of Xf_PD_^34^. Unravelling the early events of Xf_PD_ spread in the southeastern US — a geographical source of American grapevine germplasm — may allow linking crucial biogeographical events with an understanding of the historical causes of PD absence in Europe. Here, we integrate recent advances in Xf_PD_ population genetics, phylodynamic and epidemiological modelling to investigate the biogeographic events that may have precluded the introduction, establishment and spread of PD in Europe. Circumstantial evidence was compiled from several sources, including historical data on the export of American grapevines to France, a comparison of susceptibility between American grapevines and European varieties and dating of the spread of Xf_PD_ in the southeastern US. The dynamics of PD epidemic risk in Europe are accounted for the period 1870 to 2023, an extensive timeframe that provides some insight into how the climatic conditions that have historically prevented the development of PD are rapidly shifting with climate change.

## Results

### The phylloxera crisis in European vineyards preceded the spread of Xf_PD_ in the US

Between 1872 and 1900, more than 14 million cuttings of American *Vitis* species were exported from the US to France to be used as resistant rootstocks^35^. Whether or not this plant material carried Xf_PD_ infections is linked to the evolutionary history of the pathogen in the US. To shed new light on this question, we extended the phylogenetic analysis of the global Xf_PD_ population *sensu lato* to include isolates from almond and other hosts in addition to grapevine, due to their close genetic relatedness (average nucleotide identity > 99.5%; Supplementary Dataset 1 & 2). Both maximum likelihood and a preliminary Bayesian analysis without priors confirmed that the world population of Xf_PD_ (*n* = 239) is composed of two major clades, the western US clonal clade (WUCC), which includes isolates from California and Spain, and the eastern US clonal clade (EUCC), in which isolates from Taiwan are nested (Fig. 1). However, in contrast to the study by Castillo et al. 2021^24^ there is good support that the three isolates from Georgia (16M2, 16M3 and XF51 CCPMI) are basal to the whole Californian clade and are therefore included as part of the WUCC (Fig. 1a, b).

**Figure 1.**
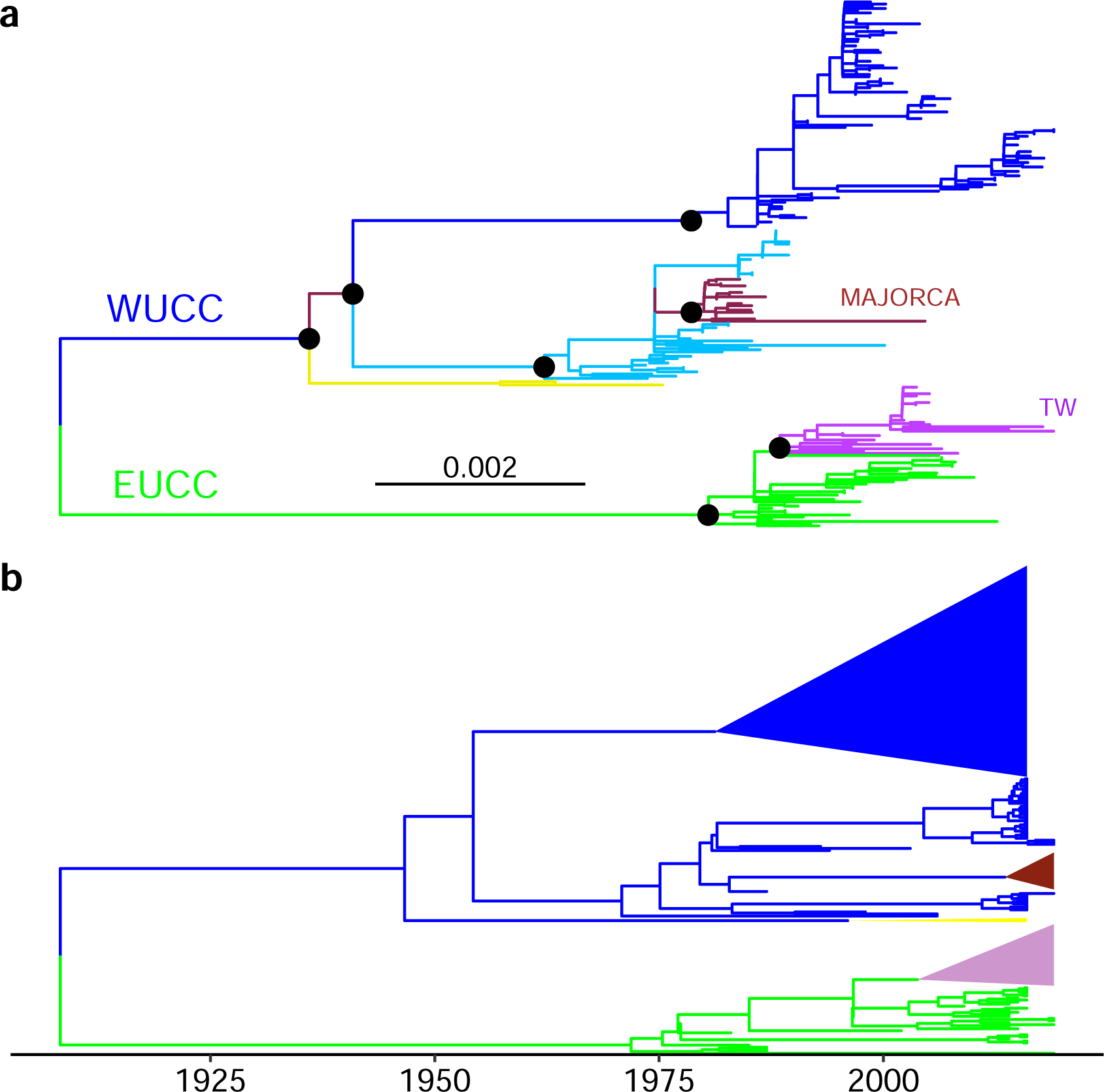
Phylogeny of worldwide Xf_PD_ isolates. (a) Midpoint rooted maximum-likelihood tree of the global Xf_PD_ population constructed using IQ-Tree and based on non-recombinant SNPs from the core genome. In yellow is the basal lineage of isolates (16M2, 16M3 and XF51 CCPMI) from the southeast of the US that form part of the western US clonal clade (WUCC). Clusters are coloured according to their geographic origin. Black asterisks indicate >90% bootstrap branch support with 1000 bootstrap replicates. (b) Maximum clade credibility using Beast without prior information. Both tree topologies reflect two old introductory events, one in California and the other in the southeastern US, followed by adaptive divergence and subsequent demographic expansion, with no subsequent migration events among demes.

While there is scientific consensus that the most likely time of introduction of Xf_PD_ into the US was around 150 years ago (∼1873)^24,25^ — a few years after phylloxera was first detected in Europe— all previously published dated phylogenies contain large uncertainties in their confidence intervals^19^. To trace the time of the most recent common ancestor (tMRCA) of the global Xf_PD_ population, we used a tip-dating strategy and a strict-calibrated molecular clock to assign dates to internal nodes of each geographic clade and subclade that showed sufficient temporal signal, such as the Taiwan and southeastern US subclades (Supplementary Fig. 1). This information was then used to estimate internal node dates and their high posterior densities (HPD) by including them as priors in the Bayesian analysis for the whole tree.

Next, we took advantage of a previous study on the epidemiology of the almond leaf scorch outbreak in Majorca to estimate the tMRCA of the subclade using a logistic-growth tree prior. A combination of a Hasegawa-Kishino-Yano (HKY) substitution model along a strict molecular clock resulted in the best-fitting model with a logistic growth rate coefficient (*k* = 0.65) very similar to that estimated for the almond leaf scorch epidemic (*k*∼ 0.6)^21^. In the extended phylogeny including isolates from *Rhamnus alaternus*, grapevines and almond trees, Xf_PD_ isolates from Majorca (*n* = 18) formed a monophyletic subclade nested within the WUCC, with the tMRCA around 2005 [95% HPD; 1966-2015] (Supplementary Fig. 2). All isolates from Taiwan (*n* = 31) clustered within the monophyletic EUCC clade (Fig. 1 and Fig. 3) and most likely originated from a single introduction event from southeastern US with the tMRCA estimated to be before 2001 [95% HPD; 1997-2002] (Supplementary Table 1). A HKY-strict clock model combined with the Bayesian skyline coalescent model yielded the best-fitting phylogeny for the EUCC clade, with the tMRCA of the 32 isolates converging around 1956 [95% HPD; 1929-1972] (Fig. 2b).

**Figure 2.**
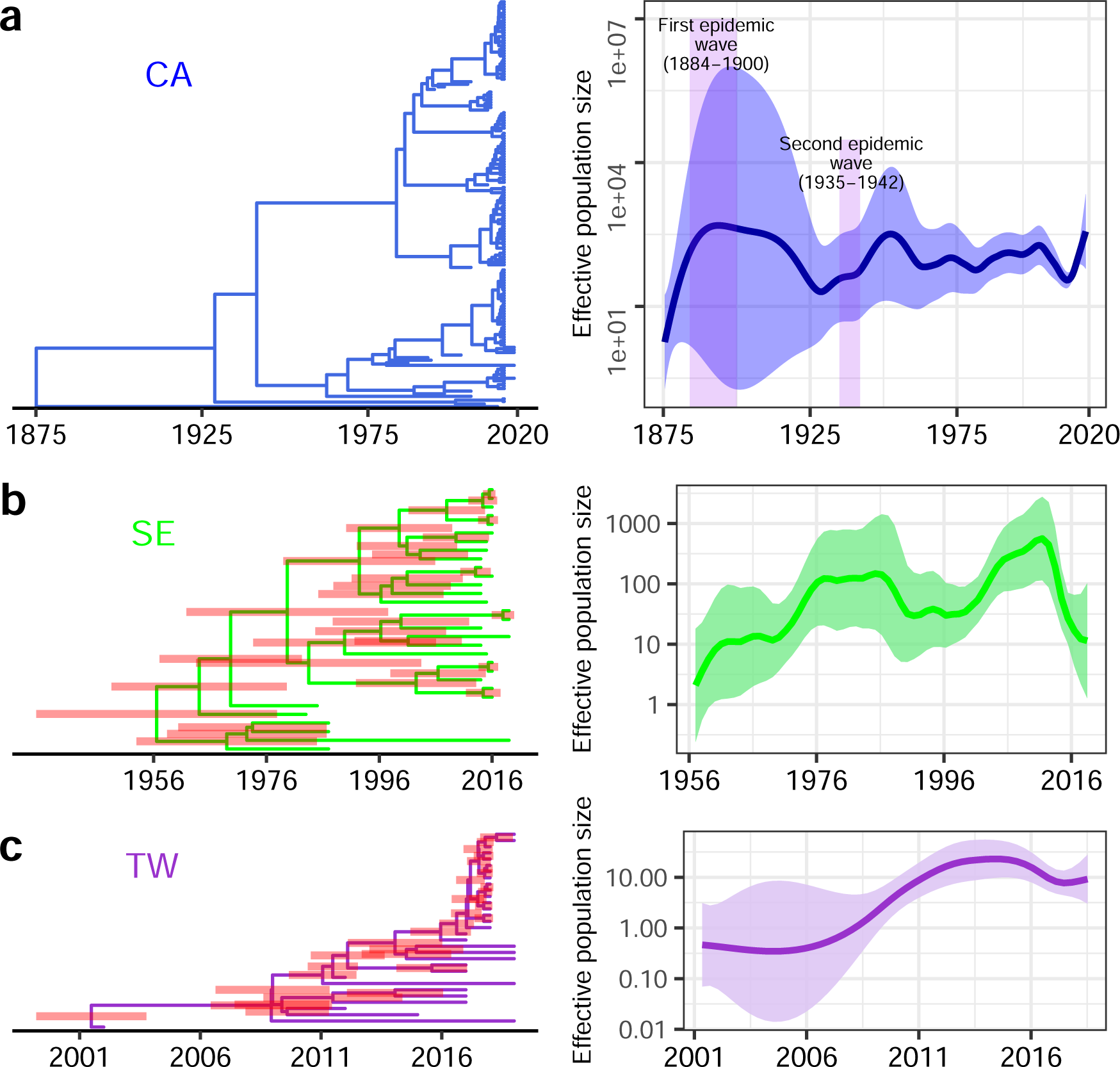
Regional expansion and emergence of Pierce’s disease epidemics. Temporal maximum clade credibility phylogenetic trees and their associated effective population size of PD epidemics in California (a), southeastern US (b) and Taiwan (c). Isolate 14B1 from Georgia was included as an outgroup in the Californian subtree to show the time of divergence between WUCC and EUCC. Node bars show 95% HPD. The estimated effective population size for the Californian clade follows the rise of the two major reported PD epidemics in the State at the end of the 19^th^ century and in the 1935-1941 period.

**Fig. 3.**
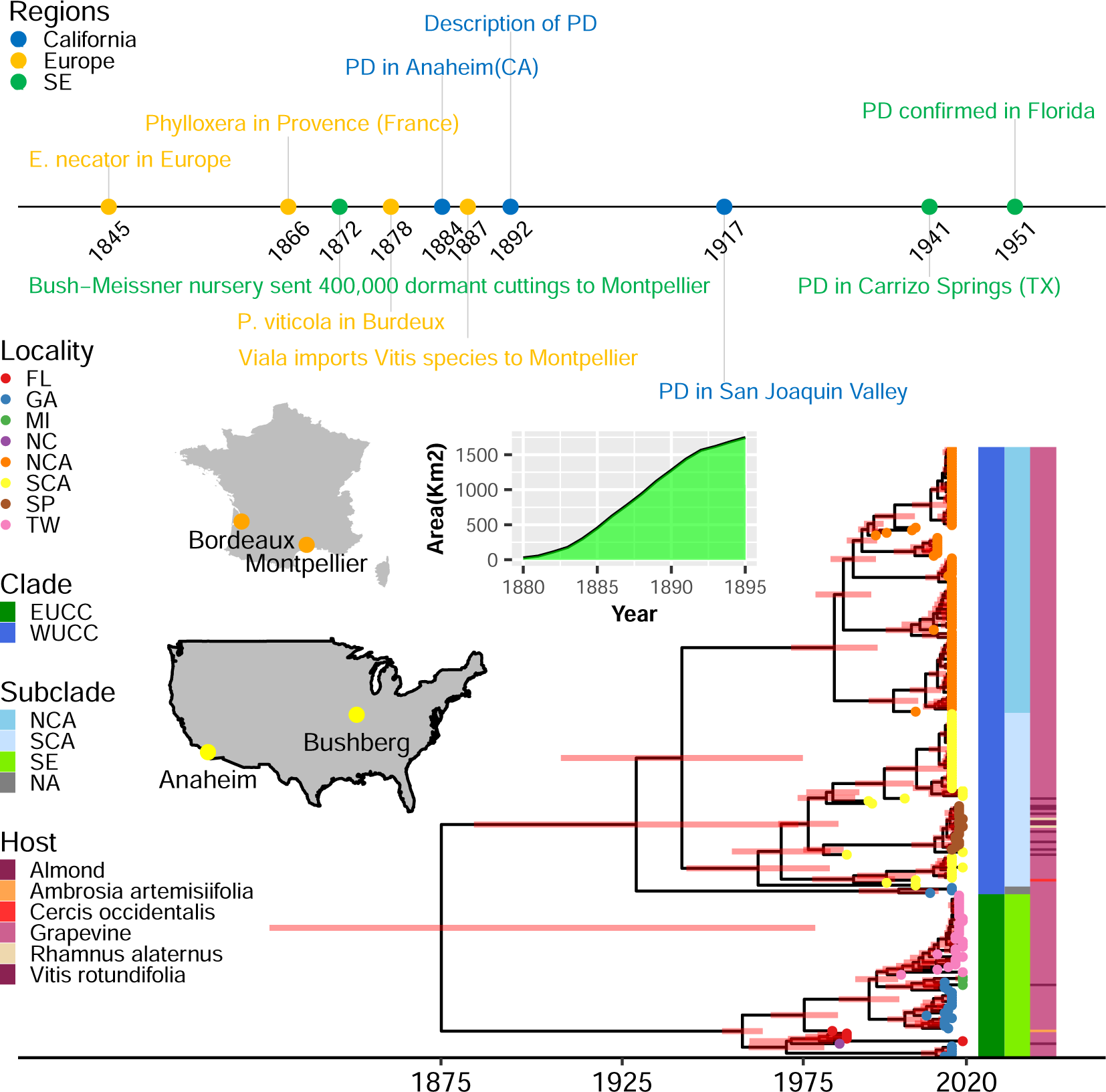
Time-calibrated maximum clade credibility phylogenetic tree of 239 Xf_PD_ genomes in the historical context of Pierce’s disease. Timeline of the major events related to vine diseases and export of plant material from US to Europe. Increase in the area planted with American rootstocks in France at the time of the phylloxera. Most American vines were grown at Bushberg in Missouri and exported to Montpellier and Bordeaux at the same time as the first description of Pierce’s disease in Anaheim, California. All these events fit quite well with the dated tree estimated using the skyline demographic model and a log-normal relaxed clock. The western US clonal clade (WUCC) and the eastern US clonal (EUCC) clades diverged at the time of the first epidemic initiated around 1875 in southern California suggesting that infected plant material was exported from California to the southeastern USA. Coloured bars show major clades, subclades and host; tree tips indicate location of Xf_PD_ isolates.

Bayesian phylogenetic analyses using the above dated nodes as strong priors confirmed the early split of the original Xf_PD_ population into two deeply branched clonal clusters, WUCC and EUCC, as previously shown only for grapevine isolates^24^. The randomizing-date test revealed significant rates of evolution in the sampling times of bacterial genome sequences (Supplementary Fig. 3). Using the best-fit model (GTR/UCLN/Coalescent-constant), we inferred a median molecular clock rate of 3.3 × 10^−7^ substitutions per site per year (95% HPD; 2.5 × 10^−7^– 4.2 × 10^−7^). Noteworthy, substitution rates were similar across models and overall slightly lower than previous estimates^19,21^. Xf_PD_ isolates from almond and other hosts were clearly interspersed with grapevine isolates within their respective geographic subclades and no migration within demes were detected (Fig. 3). The tMRCA of the northern Californian subclade was inferred to be around 1983 [95% HPD; 1969-1992] and that of the southern Californian subclade to be 1962 [95% HPD; 1937-1977]; while the time of divergence between both subclades was estimated around 1941 [95% HPD; 1901-1967] (Fig. 3). Notably, the long branches separating the WUCC and EUCC clades suggest a loss of representative samples from the Xf_PD_ ancestral populations and their replacement by a more recently adapted subpopulation^24^. In particular, the estimated tMRCA dates for both Californian subclades were substantially younger than the actual dates of the outbreaks reported by Newton Pierce in 1884 and the subsequent introduction into the Napa Valley around 1891^10^, indicating that the basal lineages to the WUCC have most likely gone extinct.

Our time-dated tree points out that the current divergence of the Californian population arose after the second major PD epidemic wave (1935-1941) and the subsequent replacement of the original Xf_PD_ population derived from the first epidemic wave (∼1882-1900)^36^ (Fig. 2a and Fig. 3). This would explain why, despite intensive and extensive sampling of Californian vineyards, no strains of this ancestral population have been detected to date^24^. It is also in line with the theoretical results expected in the periodic selection, whereby adapted ecotypes would outcompete all other strains of the former ecotype to their extinction^37^. Such genetic sweeps would be common in the case of “almost obligate” pathogens like Xf that undergo important seasonal bottlenecks. On the other hand, the time-dated tree agree with other studies in that the separation of WUCC and EUCC occurred soon after their introduction into the US^25^, as indicated in the models with highest scores in the marginal likelihood estimations (Supplementary Table 1). Although the direction of entry is unclear, the ancestral state reconstruction places 64.3% of trees in California as original location of introduction. Accordingly, the most parsimonious hypothesis is that some infected vines from the expanding population in the first epidemic wave in California were exported to the southeastern US (Supplementary Fig. S4). Of this initial population in the southeastern US, only the EUCC lineage has survived to the present day, supposedly in a process of geographic divergence and periodic selection of ecotypes more adapted to American grapevine varieties with some tolerance to PD. On the contrary, Xf_PD_ isolates from Texas, Georgia and Virginia (not included in this study) which are basal to the Californian clade would indicate a second introduction from California to the southeastern US starting with the second epidemic wave in California (1935-1941) (posterior probability = 1; Fig. 3). The current coexistence of two ecotypes today in Georgian vineyards fits again into the theoretical predictions of periodic selection whereby two independently formed ecotypes cannot compete to extinction^37^. Finally, while it is plausible to hypothesize that the Xf_PD_ strain originated from a reservoir in another host in the southeastern US and was then exported to California around 1875, currently there is no genetic clue or historical evidence to support this.

### Phylodynamics support the timeline of reported epidemics

Based on the Xf_PD_ regional genealogies, we inferred the temporal effective population size of Xf_PD_ populations from Taiwan, southeastern USA and California using Skygrowth^38^ (Fig. 2). The basic reproductive number *R_0_* estimated from the temporal phylogenetic analysis in the four geographical areas where PD occurs shows reasonable agreement with the *R_0_* estimated from the temperature-driven epidemiological model^32^ (Supplementary Fig. 5). In addition, we found a significant correlation between the growth rate of the number of infected vines inferred by Skygrowth and that risk index estimated by our epidemiological model for the southeastern US (Spearman’s rank correlation rho = 0.310, S= 27372, *p* =0.01397) and southern California CA (rho = 0.351, S= 27029, *p* =0.00476). These correlations, though weak, are remarkable given the extension of the area of study. We also observed two major demographic peaks preceded by an increase in the effective reproductive number *R*_t_ consistent with the dates of the two major PD epidemic waves reported in California between 1884-1900 and 1935-1941 (Fig. 2a) ^12^. The last epidemic wave (1935-1941 period) was much extensive and more important in the Central Valley and coincides with the emergence of the first detection of almond leaf scorch^39^. Such agreement between the root height of the WUCC tree and the PD epidemic cycles in California reinforces the accuracy of both the dated phylogeny model and the temperature-driven epidemiological model.

### PD epidemic risks in Europe from 1870 to present

Around 1872, soon after the discovery of phylloxera infestation in the Rhone Valley^40^, the first batches of imported American vines arrived in Montpellier (south-east France) and Bordeaux (Atlantic coast of France), mostly from the Bush & Son & Meissner nursery, Bushberg, Jefferson County, Missouri, US (Fig. 3). Despite the massive importation and propagation of American vines as rootstocks to combat phylloxera ^34^ (Fig. 3), there is no evidence that French vineyards or those of other European countries were affected by PD during the 19th and 20th centuries. To investigate whether this absence of PD could be attributed to unsuitable climatic conditions, we assessed whether Xf_PD_ could become established and subsequently spread among vines if we forced the introduction of infected plants. To do this, we ran our epidemiological model to retroactively determine the epidemic risk of PD in Europe^32^ using historical daily temperature projections from the Coupled Model Intercomparison Project (CMIP5) fifth phase experiment from 1850 to the present, and assuming a similar ecological range for the vector, *Philaenus spumarius*, during this period. We found negative risk indices from 1850 to 1980 in most continental France (> 98% of the territory), the only exceptions being the Provence-Alpes-Côte d’Azur (PACA) region and the island of Corsica. In the PACA region, the risk indexes were mostly in the transitional risk category (-0.1 > *r* < 0.1) between 1872 and 1920 and in the very low or low risk category from 1921 onward. In contrast, the risk index remained in the low to moderate category in Corsica (Supplementary Movie 1).

Due to the low spatial resolution (∼100 x 100 km grid) of the CMIP5 models, the epidemic risk might be underestimated at the local scale in some hotter coastal areas and river valleys of the continent. To overcome this inconvenience, we searched for the relationship between the PD risk index and annual temperature anomalies (*ΔT*) for Montpellier, Bordeaux and the PACA region and the risk index derived from our model using ERA5-Land data (∼10 x 10 km^2^ grid) from 1950 to the present. A strong non-linear correlation was observed, allowing a clear separation of *ΔT* threshold, below which the risk index would be expected to be negative (Fig. 4). Extending *ΔT* backward in time to 1850— data of temperature anomalies with coordinates are well represented from 1850 onwards —, we found that risk indexes were negative for Bordeaux from 1850 to the present, whereas risk indices were also negative (*ΔT < −*0.75) in Montpellier from 1850 to 1980, with a short positive 4-y period between 1949 and 1953 and an exponential increase in the risk index since the 1990s (Fig. 4). In the PACA region risk indexes were positive most of the time, though there were continuous periods of negative risk (*r* < 0) from 1887 to1896 and from 1902 to 1926 (Fig. 4), which were long enough to eliminate any incipient outbreak. Furthermore, Europe underwent the largest period of negative summer temperature anomalies due to volcanism forcing in the second quarter of the 19th century, with the highest peak in 1902^41^. The effect of these cool summers is reflected in the lower accumulation of growing degree days in our model when using the summer thermal anomaly instead of the annual thermal anomaly to calibrate the risk index. Overall, the risk index was reduced even further correctly capturing the model these exceptional events (Supplementary Fig. 6).

**Figure 4.**
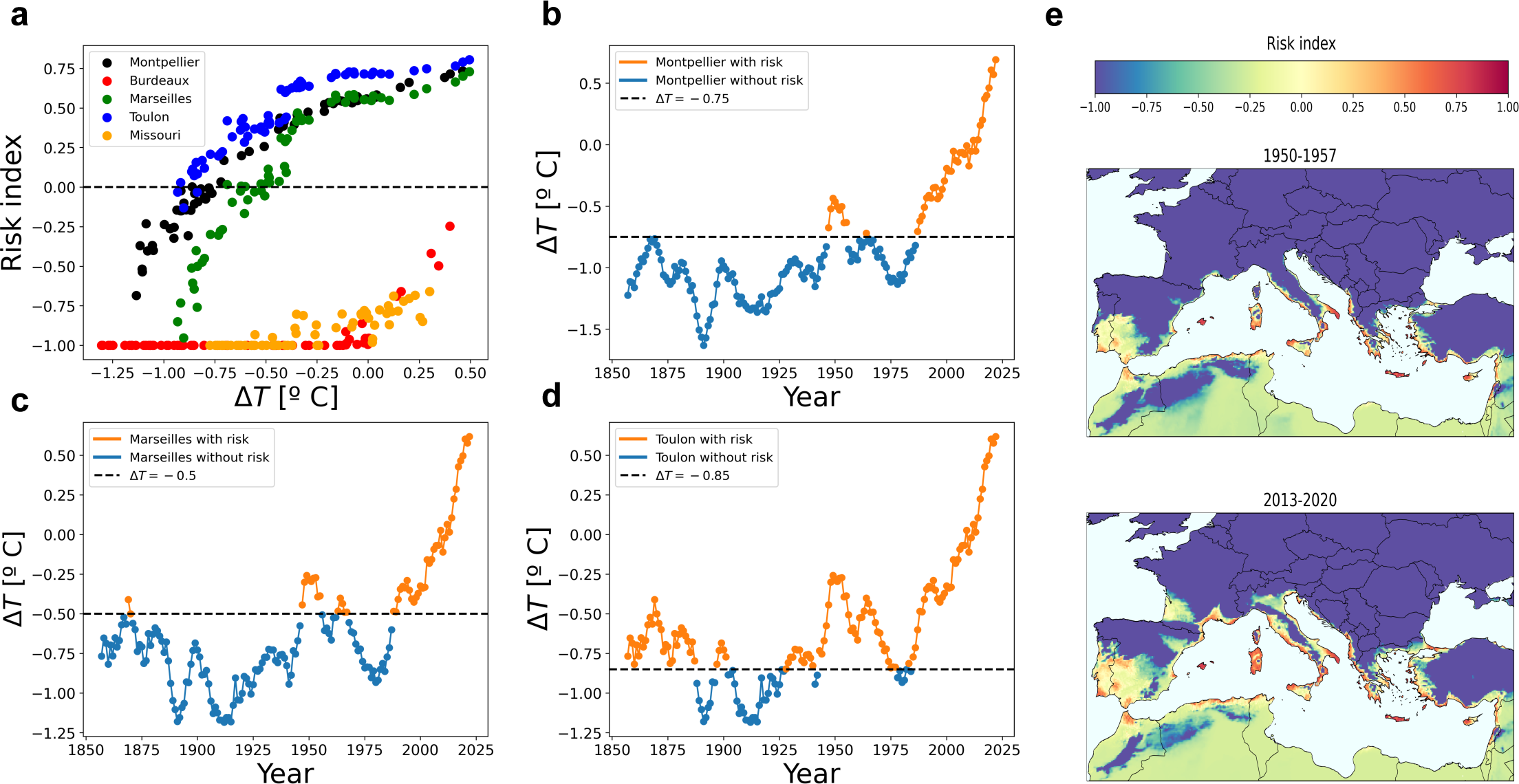
Potential PD epidemics in Europe through time. (a) Relationship between risk index and mean annual temperature anomalies for the French cities of Montpellier, Bordeaux, Marseille and Toulon and the American city of St Luis in Missouri. Data on temperature anomalies with respect to the 1991-2020 average for all locations between 1956 and 2023 were retrieved from National Oceanic and Atmospheric Administration (NOAA). Risk indices were estimated using hourly temperature data from ERA5-Land applied to the model for the same period. (b) Time series from 1850 to the present showing temperature anomalies below and above the epidemic risk threshold for Montpellier (b), Marseille (c) and Toulon (d) derived from the relationship in (a). The epidemic risk is negative throughout the period 1850-2023 in Bordeaux (not shown). (e) Snapshots showing the evolution of Pierce’s disease risk maps for Europe from 1950 to 1957 and 2013-2020. The risk index denotes the potential exponential growth of the epidemic; -0.1 to 0.1 transition risk, 0.1-0.33 low risk; 0.33-0.66 moderate risk and > 0.66 high risk. For more details see^32^.

Other evidences also argue against the possible introduction of Xf_PD_ from the US during the European phylloxera crisis. It is known that phylloxera, powdery mildew and downy mildew were introduced from the continent to the Mediterranean islands (Sicily, Crete, Sardinia, Balearic Islands, etc.) with infected vines at the end of the 19th century^42^. However, none of these plants carried X_fPD_ infections and PD, despite epidemics could have occurred (*r* > 0) on Mediterranean islands with little variation from 1850 to the present (Supplementary Movie 1 & Supplementary Fig. 7). This circumstance would reject the likelihood of long-lasting latent Xf_PD_ infections from a hypothetical introduction from American vines and subsequent disease development after exposure to suitable climatic conditions. Risk map projections using more reliable modern temperature data from 1950 to the present confirm moderate PD risks mainly in the Mediterranean islands and to a lesser extent locally in some continental Mediterranean coastal areas (Fig. 4e & Supplementary Movie 2). In addition, our model reveals temporal trends and ongoing epidemiological processes not previously identified by correlative SDMs. Since the 1990s, there have been large shifts in the extent of PD risk zones and indices (i.e. with potential exponential growth of incidence above 0; *r* > 0) (Supplementary Movie 2), mainly within transition and low-risk zones in Europe associated with a progressive increase in mean annual temperatures (Fig. 4e). Conversely, Mediterranean islands with the highest risk indexes showed low risk variation (s.e. = ± 0.1) over the same period (Supplementary Fig. 8). Over the last two decades, the largest increases from non-risk to transitional and/or low-risk zones have occurred in important wine-growing areas located in the Douro, Garonne, Rhone and Po river basins (Fig 4 & Supplementary Movie 2). These results are in fully agreement with the regional trends of climate change for Europe^43^, indicating that the model is sensitive enough to capture regional differences in climate velocity shifts in the risk index.

### Risk of PD epidemics in American vines

Just as temperatures in the late 19th century were probably unsuitable for the development of PD in most of continental Europe according to our model, temperatures in the American nurseries of origin (Supplementary Movie 1) were also likely unsuitable. Nearly all American wild vine germplasm exported to France to combat phylloxera was grown in a nursery at Bushberg, Missouri in the Midwest of the USA^40^ (Fig. 3). This location (38.268745, -90.482696) is currently well north of the *r* = 0 boundary in present risk maps, and even further if the risk index is extrapolated to the end of the 19th century, when winter temperatures would be too low for chronic infection (Supplementary Fig. 11). This can be observed in the Supplementary Fig. 12b, which shows the contractions of the PD risk area based on the thermal anomalies collected in the period (1957-2023). In North America, PD risk areas are significantly reduced in years with negative annual temperature anomalies, while in continental Europe the risk is almost zero in years with negative anomalies (Supplementary Fig. 12a). Taken together, climatic conditions were most likely unsuitable for PD development both in the nurseries of origin in the American Midwest and in France, where rootstocks were propagated and distributed throughout the country^42^.

American vine species and hybrids generally exhibit greater tolerance or/and resistance responses to Xf_PD_ than European *V. vinifera* varieties both in the field and inoculation experiments^14,34^. Consistent with this, we found that rootstocks derived from American vine hybrids developed significantly fewer severe PD symptoms and were less infected than rootstock/hybrids combinations with European varieties in the inoculation assays (Supplementary Fig. 9a). In addition, American rootstocks required approximately 1.3 times more accumulated modified growing-degree days (MGDD) to reach similar levels of symptom severity as European varieties in the 800-1027 *MGDD* interval (Supplementary Fig. 9b). To investigate the effect of climate on PD epidemics within wild *Vitis* populations, we tested our epidemic model by calibrating the average MGDD function from data obtained from American grapevine and hybrid inoculations. We obtained a risk map for the eastern US very similar to that calibrated with *V. vinifera*, a result expected mainly because summer temperatures are not a limiting factor for PD in this area (Supplementary Fig. 11 & Supplementary Movie 3). Nonetheless, lower rates of symptom development of American *Vitis* species could influence the survival of infections during winter. Even without considering the possible effect of winter curing, the limits of *r* > 0 in the PD risk map suggest that only populations of American wild vines inhabiting the coastal plains of the Gulf of Mexico (Supplementary Movie 3) could carry Xf_PD_ infections, and that plants obtained from US nurseries would have been free of Xf_PD_.

## Discussion

Our work provides new clues to interpret why PD has surprisingly not become established in continental Europe, albeit the massive importation of American vines during the 19^th^ century ^2–4^. To answer this question, we have assessed the risk of developing PD outbreaks over time conditioned by the climatic suitability of European wine-producing regions in the event of the pathogen’s introduction (*cf*. ^31^). We have also unravelled the chronology of the possible introduction and spread of Xf_PD_ in America and its subsequent migration to other parts of the world. This approach provides a robust spatiotemporal framework to address potential epidemiological, environmental and historical factors that would explain why European vineyards have escaped PD.

The historical avoidance of PD in continental Europe could be attributed in decreasing order of robustness to four main factors: (i) the non-conductive climatic conditions for the establishment of Xf_PD_ and/or its main vector in the main European wine-producing areas (Fig. 4); (ii) the importation of American grapevines from the eastern USA to Europe to combat phylloxera, which likely preceded the introduction and spread of Xf_PD_ in the southeastern US (Fig. 3); (iii) the lower susceptibility and thus bacterial load of American vines and their hybrids compared to European *V. vinifera* varieties (Supplementary Fig. 9); and (iv) the unsuitable climate for PD in the nurseries of origin (US) at the time of export to France (Supplementary Fig. 11). Each of these factors alone —if accurate— could account for the exclusion of PD in the last quarter of the 19th century. Even within the margins of temporal uncertainty of the dated phylogeny and epidemiological model, the combined likelihood of Xf_PD_ introduction, establishment and spread should be viewed as very unlikely.

Climate exclusion between 1870 and 1980 is strongly supported in our model. We highlight that the thermal dependence in the probability of developing PD symptoms was parameterized from inoculation experiments involving 36 *V. vinifera* varieties, the reliability of the model with the vector is quite high and predicts the current distribution of Xf subspecies *fastidiosa* in Europe and very likely other subspecies as well^32^. PD risk maps based on reanalysis of ERA5-Land climate data from 1950 to the present, though initially built exclusively for Xf_PD_, also explain in detail the recent developments of Xf-induced diseases in Europe (Fig. 4). This is well illustrated by the evolution of PD risk in the south-eastern Mediterranean of France from 1956 to the present (Supplementary Movie 4) and Portugal (Supplementary Movie 5), showing how the risk of establishment in these regions has only recently emerged.

Certainly, the non-interspersion of isolates from different countries within major phylogenetic clades suggests that phytosanitary controls on the international movement of grapevine germplasm have contributed to containing the spread of PD. To our knowledge, there has been no successful introduction of Xf_PD_ into Europe via infected grapevines — the only known well-established population on the island of Majorca was probably introduced with infected almond plant material from California^21^. Although these facts would indicate effective controls, our PD risk maps strongly suggest that climate rather than phytosanitary measures played a greater role in the exclusion of Xf_PD_ from European vineyards during the 20th century (Fig. 4). It seems no coincidence that the current reduced distribution of Xf in Europe is all within the moderate PD risk zones (compare Supplementary Fig. 10 and Movies 1), clearly reinforcing the predictions of our mechanistically based model. In future scenarios of +2 and +3°C annual temperature increases, we forecast that climate will no longer prevent potential outbreaks in Portugal, southern France and Italy^33^. PD prevention in these areas will therefore rely solely on surveillance and appropriate phytosanitary measures to prevent the introduction of Xf_PD._

The chronology of our dated phylogeny implies that American vines exported to France between 1875 and 1900 should have been free of Xf_PD_ infection (Fig. 3). Our Bayesian ancestral state reconstruction dates the tMRCA of the global Xf_PD_ population to 1875 [95% HPD: 1735-1941]. Although the reconstruction gives a higher probability for Californian origin, the results are still inconclusive due to the lack of sampling of Xf_PD_ both close to the root of the phylogenetic tree^44^ and in natural vegetation in the southeastern US which could help to discard its origin from this region. The most accepted hypothesis, the introduction of Xf_PD_ from Central America to southern California before 1880 and from California to the southeastern US soon after, is more in consonance with data presented here and the timeline of reports of PD and ‘degeneration disease’ in California and the southeastern US, respectively. A southeastern US origin for the Xf_PD_ metapopulation, although plausible, is not supported by genetic data and disease reports to date. Such a hypothesis stems from the observation of greater resistance/tolerance to PD in native vines compared to European *Vitis vinifera*, suggesting some form of coevolution^14^. It is also grounded in the historical disease problems of *V. vinifera* cultivated in the southeastern US^11^. Nonetheless, the “grape degeneration” of European *V. vinifera* plantations in the southeastern US was notoriously not mentioned in either the Grapevine Manual of 1883^45^ or Viala Scientific Mission Report^35^ where other grapevine diseases and pests were described in detail. The “grape degeneration” was first mentioned in Florida around 1895. Since then, certain symptoms attributed to other pathogens showed suspicious similarities with PD^11^. Many of the historical failures of European variety plantations could be explained by phylloxera alone, but also by the action of a large number of other indigenous pathogens and pests (downy mildew, powdery mildew, anthracnose, black rot, nematode-borne viruses, etc.) which could easily have been confused with PD^11^. In fact, full confirmation of PD in the southeastern USA by Californian plant pathologists did not occur until 1951^46^. On the other hand, a recent study has shown that American grapevine species from the arid regions of the western US and Mexico, where Xf_PD_ is not currently found, are more resistant to PD than vine species from the southeastern US and Mexico. These findings do not support the endemic hypothesis for the *fastidiosa* subspecies^34^. However, it cannot be ruled out that in a warmer past the distribution of the *fastidiosa* subspecies would have reached more northerly latitudes before the Pleistocene Last Glacial Maximum 18,000 years ago. The hypothesis that Xf_PD_ is a Pleistocene extinct paleoendemism recolonising the southeastern US could reconcile phylogenetic evidence for an invasive pathogen of low virulence to native vines. It has also been suggested that there may have been two independent introductions, one to California and the other to the southeast US from a source population in Central America^24^. Although also possible, this hypothesis would require two introductions from the same source population and at least two independent unlikely stochastic events involving a host jump in the initial transmission chains.

In our inoculation trials, American rootstock hybrids required on average 1.3 times more cumulative *MGDD* to develop similar levels of PD symptoms as European *V. vinifera* varieties (Supplementary Fig. 9). This lower susceptibility generally implies lower bacterial loads in infected vines, delays in symptom development, lower plant-to-plant transmission and a greater chance of winter curing at equivalent temporal settings (Supplementary Fig. 9). In our epidemiological model, Xf_PD_ transmission within natural populations of American *Vitis* spp. would be highly unlikely northward beyond the coastal plains of the Gulf of Mexico. Although these rough predictions of Xf_PD_ transmission in wild American *Vitis* populations have not yet been validated in the field, there is some circumstantial evidence to suggest that transmission is not occurring. We searched the most recent EFSA host database^47^ for confirmation of infection in wild American *Vitis* in its native habitat. Surprisingly, the only rare records are of wild vines in their natural habitat growing close to vineyards^48^ or of explants grown in nurseries and exposed under natural conditions, but not of infected plants in their natural habitat. Furthermore, in his visit to California in 1894, the prominent viticulture breeder T. V. Munsen commented to N. Pierce that he had never seen symptom like those in American native vines. This lack of infection in natural populations of wild *Vitis* spp. is also suggested by the non-detection of Xf in the National Clonal Germplasm Repository (NCGR) field genebank in Davis, CA, which contains more than 68 samples of grapevine species from the southeastern US, in which Xf_PD_ DNA was not detected by PCR analysis^49^.

Our study inevitably leads to the question of why there is no continuity in the colonisation of Xf_PD_ from Central America to the coastal plains of the Gulf of Mexico. Current climatic conditions are highly suitable for Xf_PD_ throughout its range (Supplementary Movie. 3), so explanations for ecological barriers to invasion should be sought under paleoclimatic conditions. One tantalising hypothesis, based on our model simulations is that the Pleistocene Last Glacial Maximum, 18,000 years ago, would have led to the extinction of Xf_PD_ in most of North America and created a large area of arid conditions in northern and central Mexico, unfavourable for vectors. Palaeovegetation reconstructions based on the study of palynological strata suggest an important arid zone separating the tropical forests of Central America from the warm temperate forests of the coastal plains of the Gulf of Mexico in the USA^50^. Consistent with this hypothesis, recent studies have confirmed that PD is not common in the vineyards of Mexico, but where it has been found, it has been caused by the Xf_PD_ lineage in northern Mexico and by a very different lineage, phylogenetically closer to isolates found in Costa Rica, in the vineyards of Queretaro in southern Mexico^18^.

We conclude that the lack of spatiotemporal concurrence of factors leading to the establishment of PD would explain its historical absence in continental Europe. However, rising temperatures since the 1990s are expanding PD risk zones from the coast to inland areas, increasing the possibility of outbreaks in areas that were highly unlikely before 1990. By combining a dated phylogeny of the worldwide Xf_PD_ population with PD risk modelling, we can better discriminate between temporally overlapping events and thus focus solely on the effect of climate on PD risk. Our broad spatiotemporal analysis of PD risk sheds light on the direction and intensity of climate change on the likelihood of PD outbreaks in southern Europe. Although the potential economic impact has not been addressed, the evolution of the risk index in the main European wine-producing regions provides a measure of the potential epidemic severity. An important finding of the study is the need for active surveillance in southern Europe, where the risk of PD will increase from the current very low or low risk indices to moderate risk in the > +2° scenarios^33^. Early detection of Xf_PD,_ when the potential exponential growth of PD is low, enables eradication measures that would otherwise be unfeasible when risk indices reach moderate or high levels. In February 2024, at the very last review of the manuscript, ST1 was detected in Puglia, Italy. This shows the robustness of our conclusions.

## Material and Methods

### Phylogenetic analysis of the Xf_PD_

A total of 242 *X. fastidiosa* subsp. *fastidiosa* genomes were included in this study. For Xf_PD_ phylogenetic analysis, we selected 239 isolates from different hosts sharing >99.5 % average nucleotide identity scores in the sequences of the complete genome when using JSpeciesWS^51^ (see Supplementary Dataset 1& 2). Gff files of coding sequences from previously annotated assembled genomes in the work of Castillo et al. ^24^(*n* = 182) were kindly provided by the authors. The other assembled genomes were retrieved from the NCBI nucleotide database (*n* = 19) or de novo assembled from SRAs (*n* = 41) using SPAdes v3.14.1^52^ and evaluated with QUAST v4.5^53^. Xf genome contigs in multiFASTA format were individually reordered and aligned to the Temecula1 assembly (GCA_000007245.1) reference genome using Mauve’s contig mover function^54^. The Prokka pipeline was used to individually annotate the assembled and reordered genomes^55^. A core genome alignment of the PD-causing isolates plus the three Costa Rican isolates (non-PD) and one isolate collated from a coffee plant from Mexico (CFPB8073) was created using the -e (codon aware multisequence alignment of core genes) and -n (fast nucleotide alignment) flags in Roary^56^. Ancestral genotypes of the *fastidiosa* subspecies were included in the outgroup to discard genes of the Xf_PD_ lineage under strong genetic selection that could interfere with the dated tree analysis. The resulting core genome alignment was then used to build a maximum likelihood (ML) tree with RAxML^57^ (Supplementary Fig. S13). To avoid interferes in the evolutionary inference, Gubbins v.2.2.0^58^ was utilized to generate recombination-masked full genome length and SNP-only alignments using default parameters and a minimum of three base substitutions required to identify recombination. Maximum likelihood phylogenies were obtained from the recombination-purged alignments using IQ-tree v.1.6.7.2 ^59^ under the determined best-fit nucleotide substitution model (GTR+F+G4, as determined by ModelFinder^60^) and ultrafast bootstrapping of 1000 replicates. Trees were visualized and annotated with ggtree^61^ Core genome SNP-only alignments containing 665 variable sites generated as out-put file in Gubbins were used to estimate the substitution rate, emergence times and divergence dates using Bayesian approaches as implemented in BEAST v1.10 ^62^. BEAUti xml files were manually modified to specify the number of invariant sites (A: 344747; C: 380496; G: 419538; T: 373243) in the core genome. All possible combinations (*n* = 12) of two substitution models (GTR + HKY), molecular clocks (strict, uncorrelated relaxed exponential/lognormal molecular clock) and tree demographic models (constant size coalescent, exponential population growth and Bayesian skyline coalescent) were compared using Bayes factors based on marginal likelihood estimations through path sampling and stepping-stone calculations in BEAST^62^. Each analysis consisted preliminary of two independent runs of 10 million generations sampled in every 10,000 states and then selected the five best models in the log BF for two independent runs of 100 million generations. Tree calibration was performed by assigning the isolation date to each tip of the tree (year). Temporal signal of the heterochronous sequence data was explored by performing a linear regression analysis of root-to-tip distances versus sampling time using Temptest 1.5 ^63^. A date randomization test (DTR) was performed using TipDatingBeast^64^ to further assess the genetic and temporal signal present in a dataset. MCMC convergence was assessed in Tracer v1.7.1^65^, ensuring effective sample size values higher than 200. Log-Combiner v1.10.4^62^ was used to discard 10% of the states as burn-in and to combine the trees from the two independent runs. A maximum clade credibility tree from the combined trees was obtained using TreeAnnotator v1.10.4^62^. Finally, an ancestral reconstruction of Xf_PD_ was carried out using a discrete phylogeography analysis and setting the location of isolates (California vs. southern US) as trait in BEAST v.1.10.

### Phylodynamics

To proxy the epidemic history of PD outbreaks, we estimated the effective population size of the major clades and subclades in the time-scaled phylogenies using Skygrowth^38^. This method reconstructs trends in epidemic growth and decline from time-scaled genealogies using a non-parametric autoregressive model on the growth rate as a prior for effective population size. For both Taiwan and the southeastern US, we used the best-fitting phylogeny tree obtained with a GTR and HKY-strict clock models combined with the Bayesian skyline coalescent model, respectively (Supplementary Table1). The best-fitting time-scaled phylogenetic model for Xf_PD_ was used to estimate the epidemic history for the Californian clade. All tips not belonging to the Californian clade were removed from the tree, applying the *to drop* function in *ape*, except for isolate 14B1 from Georgia used as an outgroup. The basic reproduction number *R*_0_ for each outbreak was derived from the start of the epidemic (x= tMRCA) in the plots of the reproduction *R*_e_ over time. Because the risk index of our epidemiological model somehow measures the number of infected plants (see below), it could be expected a relationship between the growth rate and the risk index of PD epidemics over time. For this, we tested a Spearman’s rank correlation rho between both variables using R. 3.5.2 version software.

### Inoculation trials on rootstocks

We tested whether rootstocks differed in their susceptibility in relation to scions (grafted onto rootstocks). The main rootstocks currently used in Europe are the result of breeding efforts to find resistance to phylloxera among native American Vitis species in the late 19th century. Rootstock R110 (Ritcher 110) is the result of a crossbreeding between *Vitis berlandieri* cv. Rességuier n°2 and *Vitis rupestris* cv. Martin; P1103 (Paulsen 1103) is the result of the crossing *V. berlandieri* cv. Rességuier n°2 and *V. rupestris* cv. Lot; 41B is a cross between *V. berlandieri* x *V. vinifera*; R140 *V. rupestris* x *V. berlandieri*; 196-17 is a cross between 1203 Couderc (*V. vinifera*-*V. rupestris*) and *Vitis riparia* cv. Gloire de Montpellier. Maps showing the distribution of main native American species are available in Supplementary figure 11. One-year-old rootstocks were supplied by a nursery in mainland Spain (Viveros Villanueva Vides, SL) and grown in 20-L plastic pots with a standard potting mix. Seven rootstock combinations were used in the inoculation tests (Supplementary Table 2). Potted plants were randomly spaced in rows of 12 plants along an insect-proof tunnel exposed to ambient temperature and watered daily to field capacity, fertilised fortnightly with a slow-release fertiliser and treated with insecticides and fungicides as needed until the end of the experiment. Two weeks before the onset of the inoculation test, leaf samples were collected from all plants and tested for the presence of *Xf* by qPCR as described elsewhere^28^.

### Isolates and inoculation

For the inoculation experiment we used two isolates of *Xf.* subsp. *fastidiosa* (ST1), XLY 2055/17 and XYL2177/18, recovered from grapevines in the island of Majorca (Spain), whose genomes have recently been sequenced ^28,66^. Isolates were grown on BYCE medium at 28°C for 7-10 days, following EPPO protocols^67^. Cells were harvested by scraping the colonies and suspending them in 1.5 ml Eppendorf tubes each with 1 ml of phosphate-buffered saline (PBS) solution until obtaining a turbid (10^8^-10^9^ cell/ml) suspension. Plants were inoculated mechanically by pin-prick inoculation^17^ with slight modifications. A 10-μl drop of the bacterial suspension was pipetted onto the leaf axil and punctured five times with an entomological needle. Eight to nine replicates per rootstock combination were inoculated with the bacterial suspension and three to four plants per rootstock with a drop of PBS as a control. Inoculation was repeated two weeks thereafter by piercing the next leaf axil above the previously inoculated ^68^.

### Disease scoring

Disease severity was rated by counting the number of symptomatic leaves above the point of inoculation at eight weeks post-inoculation (WPI) and then biweekly until the 16th week. At 12 WPI, we tested symptomatic and asymptomatic plants for *Xf* infection using qPCR, taking the petiole of the second and fifth leaf above the point of inoculation. On the 14th week, we isolated Xf from five leaves per plant of all inoculated plants following the procedure described below. We considered plants with negative qPCR results and inability to isolate Xf as not infected. A within-subject (repeated measures) factorial design was performed to evaluate differences in disease severity over time among different rootstocks. To compare the differences in susceptibility between American *Vitis* species (rootstocks) and *V. vinifera* varieties (rootstock-scion combinations), we used previously published data of disease scoring of rootstock-scion combinations^21^. Rootstock, cultivars-rootstock and time were treated as fixed factors and plant subjects as a random effect. Statistical analyses were performed using a Generalized Linear Mixed Models (GLMM) and Generalized Linear Model (GLM) with R. 3.5.2 version software. We used the *glmer* function in the R package lme4^69^ for fitting GLMMs both for the estimation of disease incidence and severity in the inoculation assays. We modelled the response variable disease incidence with the binomial error (logit-link function) and disease severity with the Poisson error (log-link function).

### Epidemiological modelling

We employed a previously published climate-driven epidemiological model to analyse the epidemic risk for PD in Europe^32^. Briefly, the model is based on a standard SIR compartmental model of PD transmission from infected to susceptible plants, which relates the transmission and recovery rates together with the vector abundance to the basic reproductive number (*R*_0_) at the onset of an outbreak. The basic reproductive number defines the exponential growth/decrease of the infected host population. To reflect the heterogeneous distribution of the vector *Philaenus spumarius* in Europe, a spatial-dependent R_0_(j) is incorporated into the model. The epidemic risk maps were computed through simulation of the epidemiological model in which each grid cell was seeded with a single infected plant at (t_0_)L=L1. The simulation ran from the initial to the final year in periods of seven years using an annual time-step, computing the incidence. Cells for which the number of infected plants had vanished at the end of each period were seeded again. This process is repeated until the last period, in which no reseeding is performed to allow the system to relax. Finally, the risk index *r* is calculated from the final number of infected plants at grid cell j. The risk index r_j_(T) implicitly defines three differential risk zones in the maps: (1) non-risk zones where r_j_L≤ − 0.1, and the number of infected plants decreases exponentially; (2) transition areas where −0.1 < rj ≤ 0.1, and (3) an epidemic risk-zone where rj > 0.1 and PD can theoretically become established and produce an outbreak—the number of infected plants increases exponentially. Full details on the model are available in previous works^32,33^

### Climate data

Climate data between 1850 and 2023 from different sources were used to generate historical PD risk maps. Maximum and minimum daily temperature projections of each of the climate models were downloaded from the Coupled Model Intercomparison Project (CMIP5) of the fifth phase of the experiment from 1850 to the present (https://cds.climate.copernicus.eu/cdsapp#!/dataset/projections-cmip5-daily-single-levels?tab=overview). Hourly temperature values were computed using a sinusoidal extrapolation relating maximum and minimum daily temperature to hourly temperatures^33^. Of 16 climate models, we discarded those that reflected temporal and spatial inconsistencies in the risk map simulations. Eventually, seven models, ’CESM1-BGC’, ’CSIRO-Mk3-6-0’, ’NorESM1-M’, ’ACCESS1-0’ and ’CCSM4’, ’IPSL-CM5A-MR’ and ’ACCESS1-3’ were selected to average the infection probabilities of the US and European maps. The spatial resolution of all models was standardised to an average resolution of 1° using a first order remapping climate data operator implemented by the Max-Plank Institute for Meteorology. In the PD risk simulations, we assumed that the population density and distribution of the insect vector *Philaenus spumarius* in Europe did not vary in the period of study. Due to the low resolution of the CMIP5 climate data, risk maps from 1950 to the present were constructed from ERA5-Land data^70^ with a resolution of 0.1°. The data were downloaded in GRIB format with hourly temporal resolution and 0.1° spatial resolution. A Julia library^71^ was developed using the GRIB.jl package^38^ to compute the annual modified growing degree days and cold DD estimates. The risk of PD in Europe and the USA before 1950 was determined by evaluating the annual thermal anomalies and the risk of PD with the models simulated between 1950-2023 with the reanalysed ERA5-Land data. The data on thermal anomalies between 1850 and 2023 with respect to the 1901-2000 average (by cells) and with respect to the 1991-2020 average (coordinates) were obtained from the National Oceanic and Atmospheric Administration (NOAA) (https://www.ncei.noaa.gov/access/monitoring/climate-at-a-glance/global/time-series).

### Literature review and historical data

We focused on obtaining historical data on the incidence of Pierce’s disease in California and the southeastern US and on the export of plants from the eastern US to France between 1870 and 1900. Records of natural infections of wild American *Vitis* spp. were searched in the updated EFSA host database^47^ and Google Scholar. Distribution records of native American *Vitis* spp. were downloaded from the United States Department of Agriculture (https://plants.usda.gov/home). Vineyards area data were obtained from the National Agriculture Statistics Service of the USDA (https://www.nass.usda.gov/) and the International Organisation of Vine and Wine (https://www.oiv.int/).

## Acknowledgments

The authors thank M. Montesinos (Tragsa) and S. Martínez (Neiker) for carrying out the inoculations on *Vitis* rootstocks; R. Almeida (UC Berkley) and M. Donegan (UC Berkley) for gently providing the files with annotated assembled genomes; and L. de la Fuente (Auburn University) for discussion on the manuscript. E.M. received support by the European Research Executive Agency-HORIZON Research and Innovation Actions in the project “Beyond Xylella, Integrated Management Strategies for Mitigating Xylella fastidiosa impact in Europe” (Bexyl), grant number 101060593. A.G.R. and M.A.M. were supported through grant PID2021-123723OB-C22 (CYCLE) funded by MICIU/AEI/10.13039/501100011033 and by “ERDF A way of making Europe” and through grant CEX2021-001164-M (María de Maeztu Program for Units of Excellence in R&D) funded by MICIU/AEI/10.13039/501100011033.

## Author Contribution

E.M., A.G.R and M.A.M. conceptualized the project and conducted investigations; E.M. and A.G.R performed the numerical experiments and A.G.R performed the risk simulations and the analysis. E.M. wrote the manuscript with input from A.G.R and M.A.M. All the authors reviewed the manuscript.

## Data availability

List of genomes used in the study are provided in extended Database 1. Files with data of annotated genomes, multiple sequence alignments with and without recombining regions removed, as wells as the phylogenetic models built with Beauti and beast output log, ops and tree files needed to reproduce the phylogenetic analyses are available in https://figshare.com/s/39a6feef05b6adf3dc1d. Hourly temperature data from 1950 to 2023 necessary to prepare Fig. 4, Supplementary figures 6, 7 and 8 and movies 2 and 4 were taken from Copernicus ERA5-Land (namely the ’2m temperature’ field). Maximum and minimum daily temperature data from CHELSA dataset downloaded to build movie 5 are available at Zenodo, doi:10.5281/zenodo.10579689 (2024).

## Code availability

R scripts used for statistical analyses and plotting are available in

https://figshare.com/s/39a6feef05b6adf3dc1d. A library built in Julia to analyse the data outputs of ERA5-Land in GRIB format is available in this Github link https://github.com/agimenezromero/PierceDisease-GlobalRisk-Predictions. The simulation code and a small reproducible example are provided in this Github link https://github.com/agimenezromero/PierceDisease-GlobalRisk-Predictions.

## References

1. Borkar, S. G. History of Plant Pathology. (CRC Press, 2017).

2. Brewer, M. T. & Milgroom, M. G. Phylogeography and population structure of the grape powdery mildew fungus, *Erysiphe necator*, from diverse *Vitis* species. BMC evolutionary biology (2010) doi:10.1186/1471-2148-10-268.

3. Tello, J., Mammerler, R., Čajić, M. & Forneck, A. Major Outbreaks in the Nineteenth Century Shaped Grape Phylloxera Contemporary Genetic Structure in Europe. Scientific Reports (2019) doi:10.1038/s41598-019-54122-0.

4. Rouxel, M. et al. Geographic distribution of cryptic species of *Plasmopara viticola* causing downy mildew on wild and cultivated grape in Eastern North America. Phytopathology (2014) doi:10.1094/PHYTO-08-13-0225-R.

5. Hulme, P. E. Trade, transport and trouble: managing invasive species pathways in an era of globalization. Journal of applied ecology 46, 10–18 (2009).

6. Hopkins, D. L. & Purcell, A. H. *Xylella fastidiosa*: cause of Pierce’s disease of grapevine and other emergent diseases. Plant disease 86, 1056–1066 (2002).

7. Golino, D. A. et al. Regulatory Aspects of Grape Viruses and Virus Diseases: Certification, Quarantine, and Harmonization. in *Grapevine Viruses: Molecular Biology, Diagnostics and Management* (eds. Meng, B., Martelli, G. P., Golino, D. A. & Fuchs, M.) 581–598 (Springer International Publishing, Cham, 2017). doi:10.1007/978-3-319-57706-7_28.

8. Janse, J. D. & Obradovic, A. *Xylella fastidiosa*: its biology, diagnosis, control and risks. Journal of Plant Pathology S35–S48 (2010).

9. Hoddle, M. S. The potential adventive geographic range of glassy-winged sharpshooter, *Homalodisca coagulata* and the grape pathogen *Xylella fastidiosa*: implications for California and other grape growing regions of the world. Crop Protection 23, 691–699 (2004).

10. 10. Pierce, N. B. The California Vine Disease: A Preliminary Report of Investigations. USDA Division of Vegetable Pathology vol. bulletin n (1892).

11. Stoner, W. N. A Comparison between Grape Degeneration in Florida and Pierce’s Disease in California. The Florida Entomologist 35, 62–68 (1952).

12. Hewitt, W. B., Frazier, N. W., Jacob, H. E. & Freitag, J. H. Pierce’s disease of grapevines. Pierce’s disease of grapevines. 1–32 (1942).

13. Davis, M. J., Purcell, A. H. & Thomson, S. V. Pierce’s disease of grapevines: Isolation of the causal bacterium. Science (1978) doi:10.1126/science.199.4324.75.

14. Hewitt, W. B. The probable home of Pierce’s disease virus. American Journal of Enology and Viticulture 9, 94–98 (1958).

15. Schuenzel, E. L., Scally, M., Stouthamer, R. & Nunney, L. A multigene phylogenetic study of clonal diversity and divergence in North American strains of the plant pathogen *Xylella fastidiosa*. Applied and environmental microbiology 71, 3832–3839 (2005).

16. Yuan, X. et al. Multilocus sequence typing of *Xylella fastidiosa* causing Pierce’s disease and oleander leaf scorch in the United States. Phytopathology 100, 601–611 (2010).

17. Almeida, R. P. P. & Purcell, A. H. Biological Traits of *Xylella fastidiosa* Strains from Grapes and Almonds. Applied and Environmental Microbiology 69, 7447–7452 (2003).

18. Aguilar-Granados, A. et al. Genetic Diversity of *Xylella fastidiosa* in Mexican Vineyards. Plant Disease 105, 1490–1494 (2021).

19. Vanhove, M. et al. Genomic diversity and recombination among *Xylella fastidiosa* subspecies. Applied and environmental microbiology 85, e02972–18 (2019).

20. Su, C.-C. et al. Pierce’s Disease of Grapevines in Taiwan: Isolation, Cultivation and Pathogenicity of Xylella fastidiosa. Journal of Phytopathology 161, 389–396 (2013).

21. Moralejo, E. et al. Phylogenetic inference enables reconstruction of a long-overlooked outbreak of almond leaf scorch disease (*Xylella fastidiosa*) in Europe. Communications biology 3, 1–13 (2020).

22. Zecharia, N., et al. *Xylella fastidiosa* outbreak in Israel: population genetics, host range and temporal and spatial distribution analysis. Phytopathology (2022).

23. Vanhove, M., Sicard, A., Ezennia, J., Leviten, N. & Almeida, R. P. Population structure and adaptation of a bacterial pathogen in California grapevines. Environmental microbiology 22, 2625–2638 (2020).

24. Castillo, A. I. et al. Allopatric Plant Pathogen Population Divergence following Disease Emergence. Applied and Environmental Microbiology 87, e02095–20 (2021).

25. Nunney, L. et al. Population genomic analysis of a bacterial plant pathogen: novel insight into the origin of Pierce’s disease of grapevine in the US. PLoS One 5, e15488 (2010).

26. Saponari, M., Boscia, D., Nigro, F. & Martelli, G. P. Identification of DNA sequences related to *Xylella fastidiosa* in oleander, almond and olive trees exhibiting leaf scorch symptoms in Apulia (Southern Italy). Journal of Plant Pathology 95, (2013).

27. Landa, B. B. et al. Emergence of a plant pathogen in Europe associated with multiple intercontinental introductions. Applied and Environmental Microbiology 86, e01521–19 (2020).

28. Moralejo, E. et al. Insights into the epidemiology of Pierce’s disease in vineyards of Mallorca, Spain. Plant Pathology 68, 1458–1471 (2019).

29. Cornara, D. et al. Transmission of *Xylella fastidiosa* by naturally infected *Philaenus spumarius* (Hemiptera, Aphrophoridae) to different host plants. Journal of Applied Entomology 141, 80–87 (2017).

30. Godefroid, M., Cruaud, A., Streito, J.-C., Rasplus, J.-Y. & Rossi, J.-P. *Xylella fastidiosa*: climate suitability of European continent. Scientific Reports 9, 1–10 (2019).

31. Schneider, K. et al. Impact of *Xylella fastidiosa* subspecies *pauca* in European olives. Proceedings of the National Academy of Sciences of the United States of America (2020) doi:10.1073/pnas.1912206117.

32. Giménez-Romero, A. et al. Global predictions for the risk of establishment of Pierce’s disease of grapevines. Commun Biol 5, 1–13 (2022).

33. Giménez-Romero, À., Iturbide, M., Moralejo, E., Gutiérrez, J. M. & Matías, M. A. Contrasting Patterns of Pierce’s Disease Risk in European Vineyards Under Global Warming. bioRxiv 2023.07. 17.549293 (2023).

34. Riaz, S., Tenscher, A. C., Heinitz, C. C., Huerta-Acosta, K. G. & Walker, M. A. Genetic analysis reveals an east-west divide within North American *Vitis* species that mirrors their resistance to Pierce’s disease. PLOS ONE 15, e0243445 (2020).

35. Viala, P. Une Mission Viticole En Amérique. (C. Coulet, 1889).

36. Hewitt, W. B. Virus diseases of grapevines. US Dep. Agric. Yearbook (1953).

37. Selander, R. K., Musser, J. M., Caugant, D. A., Gilmour, M. N. & Whittam, T. S. Population genetics of pathogenic bacteria. Microbial pathogenesis 3, 1–7 (1987).

38. Volz, E. M. & Didelot, X. Modeling the growth and decline of pathogen effective population size provides insight into epidemic dynamics and drivers of antimicrobial resistance. Systematic Biology 67, 719–728 (2018).

39. Mircetich, S. M., Lowe, S. K., Moller, W. J. & Nyland, G. Etiology of almond leaf scorch disease and transmission of the causal agent. Phytopathology 66, 17–24 (1976).

40. Sorensen, C. W., Smith, E. H., Smith, J. & Carton, Y. Charles V. Riley, France, and Phylloxera.American Entomologist 54, 134–149 (2008).

41. Luterbacher, J., Dietrich, D., Xoplaki, E., Grosjean, M. & Wanner, H. European seasonal and annual temperature variability, trends, and extremes since 1500. Science 303, 1499–1503 (2004).

42. Gale, G. Saving the vine from phylloxera: a never ending battle. Wine: A scientific exploration 70, 91 (2002).

43. Loarie, S. R. et al. The velocity of climate change. Nature 462, 1052–1055 (2009).

44. Lemey, P., Rambaut, A., Drummond, A. J. & Suchard, M. A. Bayesian phylogeography finds its roots. PLoS computational biology 5, e1000520 (2009).

45. Bush, firm, Bushberg & Meissner), M. (1895 B. & S. &. Illustrated Descriptive Catalogue of American Grape Vines: A Grape Growers’ Manual. (RP Studley & Company, printers, 1895).

46. Hewitt, W. B., Loomis, N. H., Overcash, J. P. & Parris, G.-K. Pierce’s disease virus in Mississippi and other southern States. (1958).

47. Authority (EFSA), E. F. S., Delbianco, A., Gibin, D., Pasinato, L. & Morelli, M. Update of the *Xylella* spp. host plant database–systematic literature search up to 30 June 2021. EFSA Journal 20, e07039 (2022).

48. Morano, L. D. et al. Initial genetic analysis of *Xylella fastidiosa* in Texas. Current Microbiology 56, 346–351 (2008).

49. Stover, E., Riaz, S. & Walker, M. A. PCR screening for *Xylella fastidiosa* in grape genebank accessions collected in the southeastern United States. American journal of enology and viticulture 59, 437–439 (2008).

50. Domínguez-Vázquez, G., Osuna-Vallejo, V., Castro-López, V., Israde-Alcántara, I. & Bischoff, J. A. Changes in vegetation structure during the Pleistocene–Holocene transition in Guanajuato, central Mexico. Vegetation History and Archaeobotany 28, 81–91 (2019).

51. Richter, M., Rosselló-Móra, R., Oliver Glöckner, F. & Peplies, J. JSpeciesWS: a web server for prokaryotic species circumscription based on pairwise genome comparison. Bioinformatics 32, 929–931 (2016).

52. Bankevich, A. et al. SPAdes: a new genome assembly algorithm and its applications to single-cell sequencing. Journal of computational biology 19, 455–477 (2012).

53. Gurevich, A., Saveliev, V., Vyahhi, N. & Tesler, G. QUAST: quality assessment tool for genome assemblies. Bioinformatics 29, 1072–1075 (2013).

54. Darling, A. E., Mau, B. & Perna, N. T. Progressivemauve: Multiple genome alignment with gene gain, loss and rearrangement. PLoS ONE (2010) doi:10.1371/journal.pone.0011147.

55. Seemann, T. Prokka: rapid prokaryotic genome annotation. Bioinformatics 30, 2068–2069 (2014).

56. Page, A. J. et al. Roary: rapid large-scale prokaryote pan genome analysis. Bioinformatics 31, 3691–3693 (2015).

57. Stamatakis, A. Using RAxML to infer phylogenies. Current protocols in bioinformatics 51, 6.14.1–6.14.14 (2015).

58. Croucher, N. J. et al. Rapid phylogenetic analysis of large samples of recombinant bacterial whole genome sequences using Gubbins. Nucleic acids research 43, e15–e15 (2015).

59. Nguyen, L.-T., Schmidt, H. A., Von Haeseler, A. & Minh, B. Q. IQ-TREE: a fast and effective stochastic algorithm for estimating maximum-likelihood phylogenies. Molecular biology and evolution 32, 268–274 (2015).

60. Kalyaanamoorthy, S., Minh, B. Q., Wong, T. K., Von Haeseler, A. & Jermiin, L. S. ModelFinder: fast model selection for accurate phylogenetic estimates. Nature methods 14, 587– 589 (2017).

61. Yu, G., Smith, D. K., Zhu, H., Guan, Y. & Lam, T. T.-Y. ggtree: an R package for visualization and annotation of phylogenetic trees with their covariates and other associated data. Methods in Ecology and Evolution 8, 28–36 (2017).

62. Drummond, A. J., Suchard, M. A., Xie, D. & Rambaut, A. Bayesian phylogenetics with BEAUti and the BEAST 1.7. Molecular Biology and Evolution 29, 1969–1973 (2012).

63. Rambaut, A., Lam, T. T., Max Carvalho, L. & Pybus, O. G. Exploring the temporal structure of heterochronous sequences using TempEst (formerly Path-O-Gen). Virus evolution 2, vew007 (2016).

64. Rieux, A. & Khatchikian, C. E. TIPDATINGBEAST: An R package to assist the implementation of phylogenetic tip[dating tests using beast. Molecular Ecology Resources 17, 608–613 (2017).

65. Rambaut, A., Drummond, A. J., Xie, D., Baele, G. & Suchard, M. A. Posterior summarization in Bayesian phylogenetics using Tracer 1.7. Systematic biology 67, 901–904 (2018).

66. Gomila, M. et al. Draft Genome Resources of Two Strains of *Xylella fastidiosa* XYL1732/17 and XYL2055/17 Isolated from Mallorca Vineyards. Phytopathology 109, 222–224 (2019).

67. PM 7/24 (3) Xylella fastidiosa. EPPO Bulletin 48, 175–218 (2018).

68. Wallis, C. M., Wallingford, A. K. & Chen, J. Grapevine rootstock effects on scion sap phenolic levels, resistance to *Xylella fastidiosa* infection, and progression of Pierce’s disease. Frontiers in Plant Science 4, (2013).

69. 69. Bates, D., Mächler, M., Bolker, B. & Walker, S. Fitting linear mixed-effects models using lme4. *arXiv preprint arXiv:1406.*5823 (2014).

70. Hersbach, H. et al. The ERA5 global reanalysis. Quarterly Journal of the Royal Meteorological Society 146, 1999–2049 (2020).

71. Bezanson, J., Edelman, A., Karpinski, S. & Shah, V. B. Julia: A fresh approach to numerical computing. SIAM review 59, 65–98 (2017).

