## Supplementary Information for "Linking intercontinental biogeographic events to decipher how European vineyards escaped Pierce’s disease"

**This PDF file includes:**

- Figures S1 to S13
- Tables S1 to S2
- Legends for Movies S1 to S5
- Legends for Datasets S1

**Other supporting materials for this manuscript include the following:**

- Movies S1 to S5
- Datasets S1

### Supplementary figures

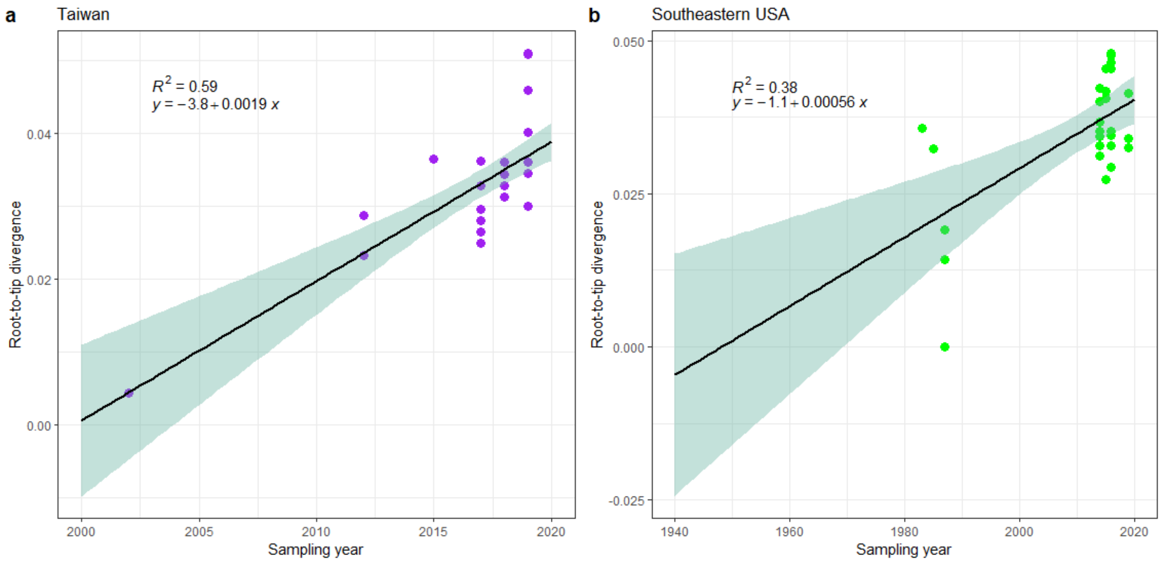


**Figure S1**. Relationship between the root-to-tip divergence and sampling dates of $Xf_{PD}$ isolates from Taiwan (a) and southeastern US (b) estimated using Temptest and plotted with ggplot2. The blue-shaded areas denote the 95% confidence interval in the regression lines.

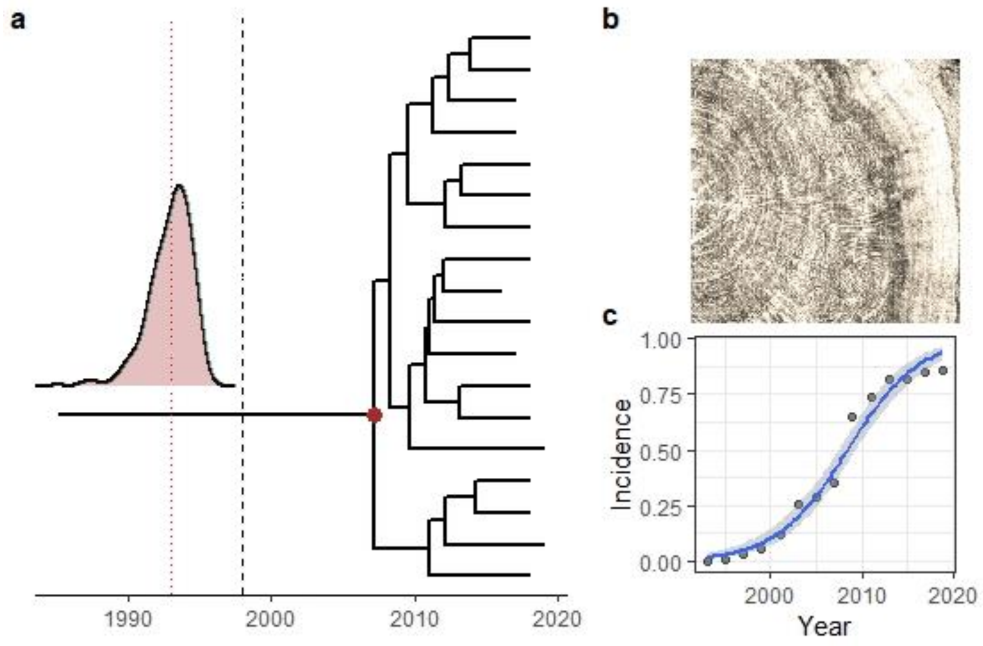


**Figure S2**. Dated phylogeny for the Majorcan $Xf_{PD}$ subclade. (a) The tMRCA estimated is based on information of Xf DNA detected on growing-rings (b) and the disease growth curve of the almond leaf scorch (c). Black dotted line indicate the minimum bound (1998) for the introduction of $Xf_{PD}$. Red dashed line the estimated time of introduction.

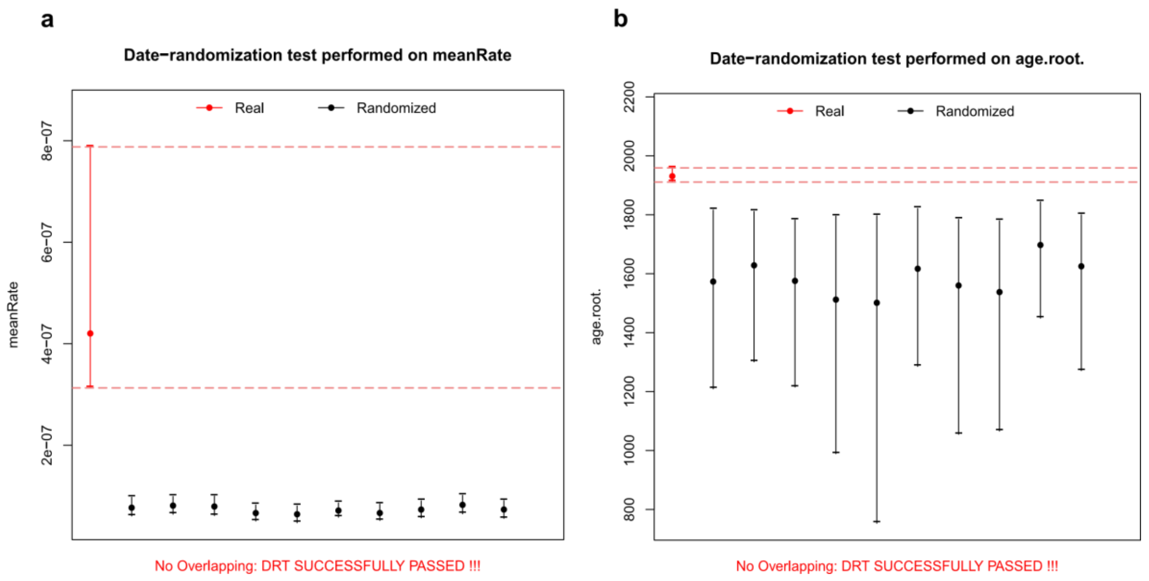


**Figure S3. Date-randomization test on $Xf_{PD}$ heterochronous samples.** (a) Temporal signal of the mean substitution rate in the time structured data shows no overlap within the 95% credible intervals with those generated with random dates as well (b) as for the root height.

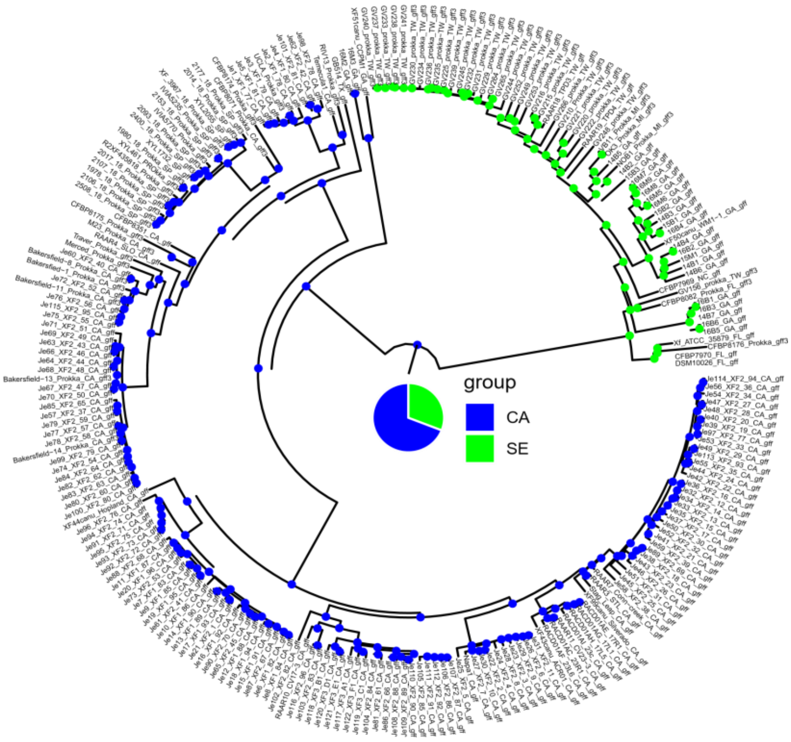


**Figure S4. Ancestral locations based on discrete trait model in Beast.** Root location probabilities of the reconstructed tree are consistent with the timeline order of the reported outbreaks in California (1884; 64.3%) and the degeneration grapevine disease in Florida around 1895. Node size are proportional to the location probability. The absence of sequences sampled close to the root can hinder the accurate estimation of $Xf_{PD}$ geographic origins.

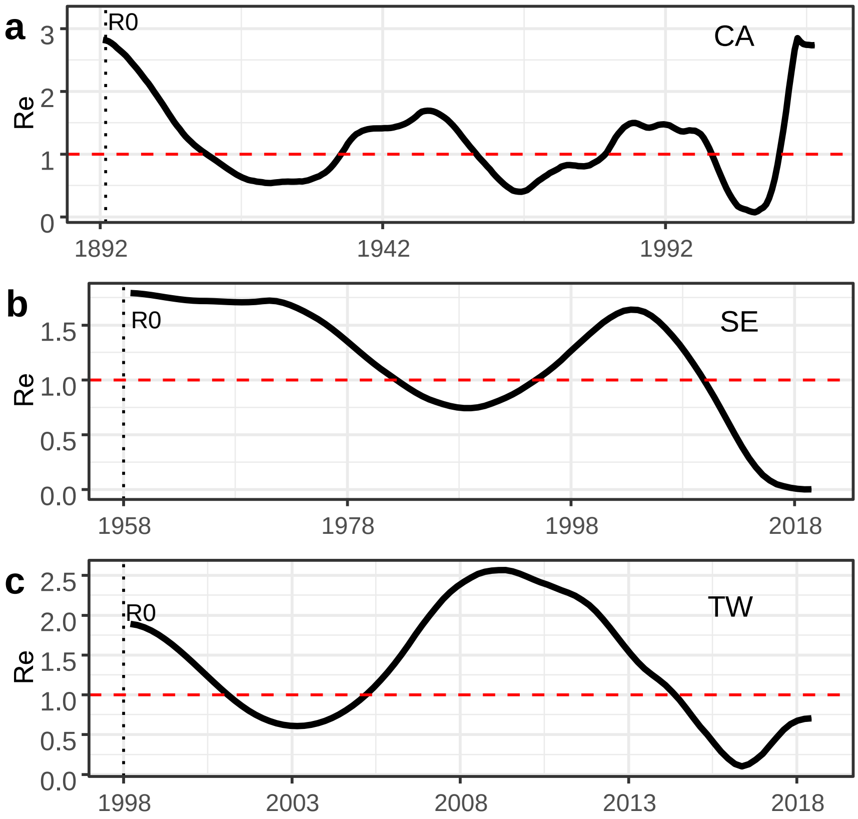


**Figure S5.** Reproductive number *Re* estimated using Skygrowth for California (a), southern US (b) and Taiwan (c). Red dashed line represents the $R_0$ threshold.

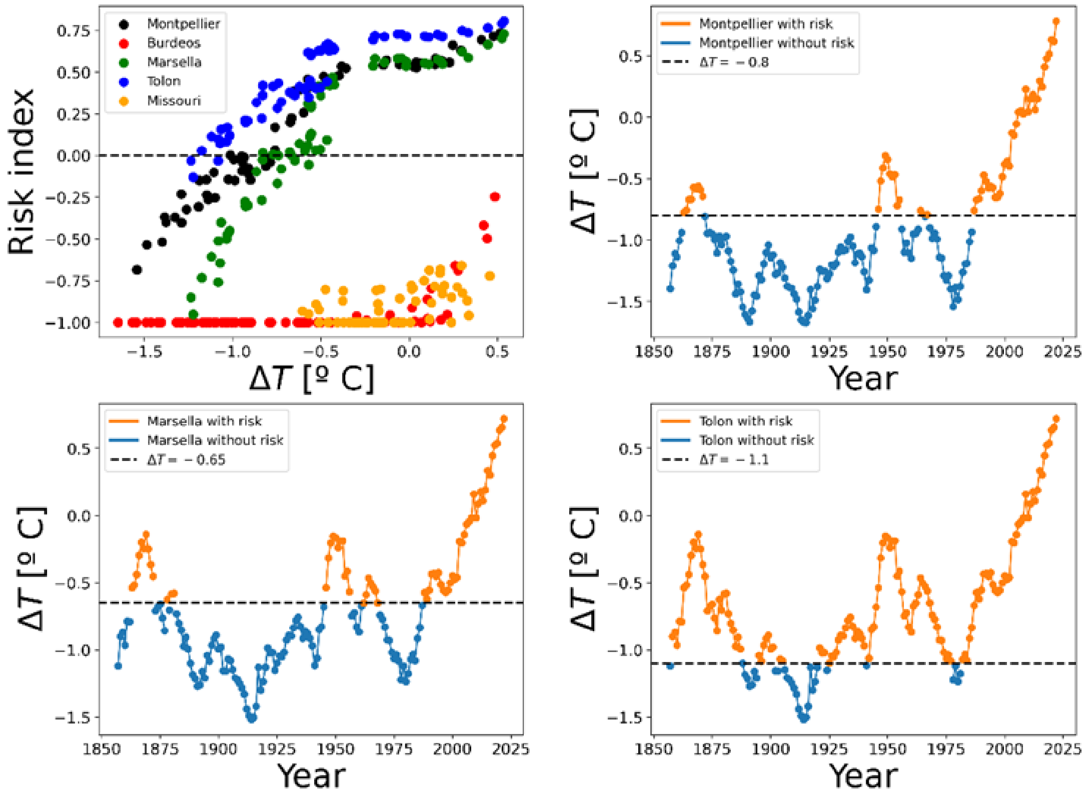


**Figure S6**. Relationship between risk index and mean summer temperature anomalies. Dash horizontal lines depict the threshold below which risk index are negative. Values below risk in blue and with risk in orange.

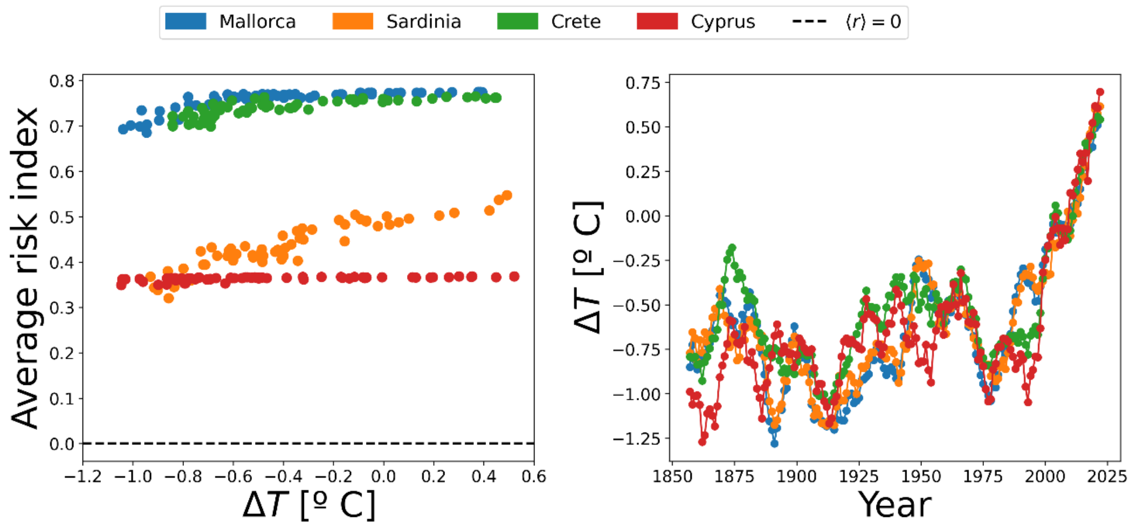


**Figure S7.** Pierce's disease risk and temperature anomalies in the Mediterranean islands. Risk indices remain quite constant independently of temperature anomalies except for Sardinia which slightly increases with positive temperature anomalies. Temperature anomalies follow similar trends in the four islands over time showing an important increase after the 1990s.

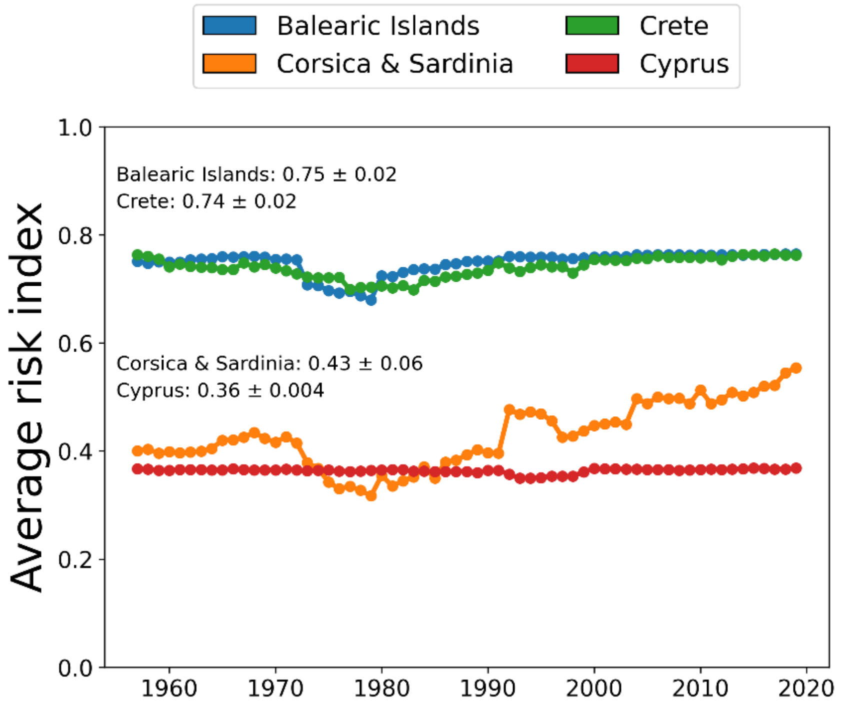


**Figure S8.** Evolution of the risk index over time in the Mediterranean Islands. The risk index remains almost constant, with little oscillation in the islands of lower latitudes and a slight upward trend in Corsica compared to continental locations as shown in Fig. S2.

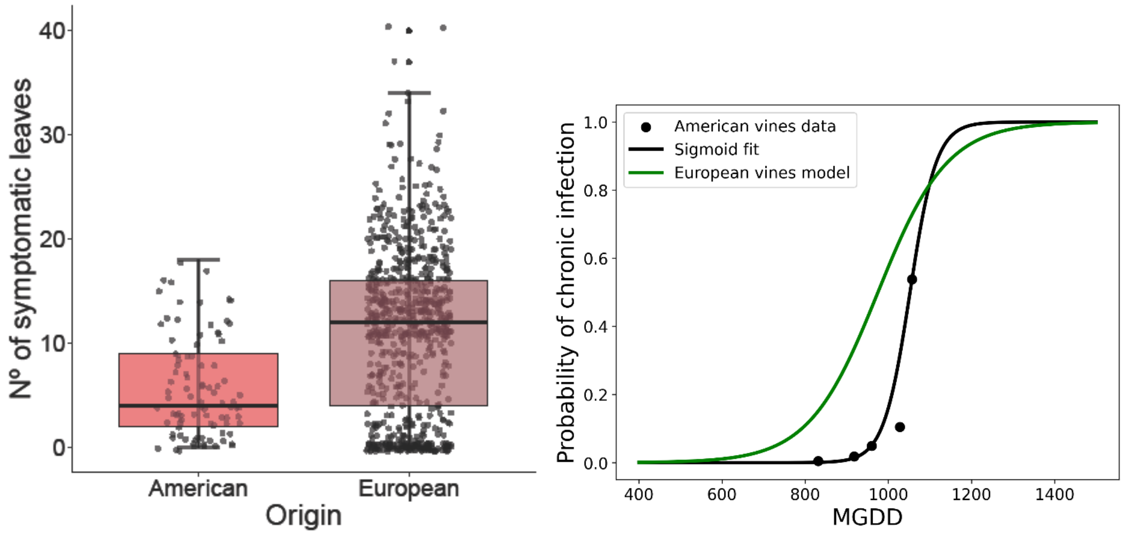


**Figure S9.** Different responses to $Xf_{PD}$ infection between European *Vitis vinifera* cultivars and American *Vitis* species and hybrids. (a) Number of symptomatic leaves shown 14 weeks after inoculation of seven rootstocks of American origin and 36 European *Vitis vinifera* varieties. (b) Modified growing degree days (MGDD) accumulated to reach an average of 5 symptomatic leaves across American vines (black line) and European *V. vinifera* varieties.

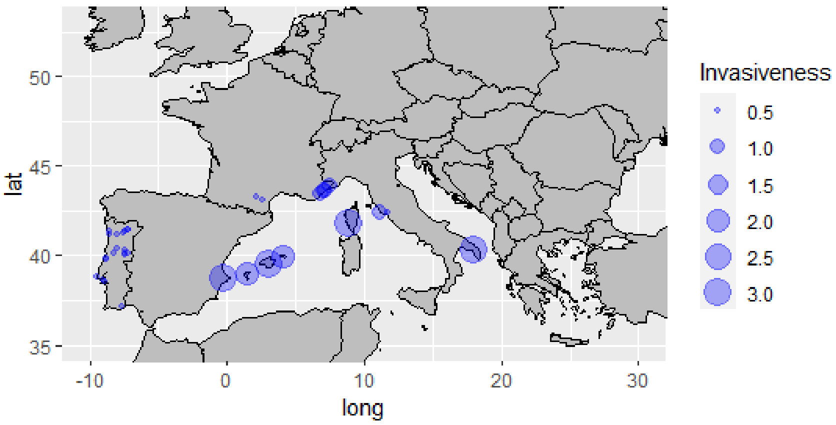


**Figure S10.** Distribution of *Xylella fastidiosa* outbreaks in Europe. The sizes of the dots are proportional to the level of establishment and invasiveness somehow reflecting the climatic suitability over time. Recent outbreaks in Occitanie (France), Portugal and Lazio (Italy) are the likely consequence of the warming in the last decade.

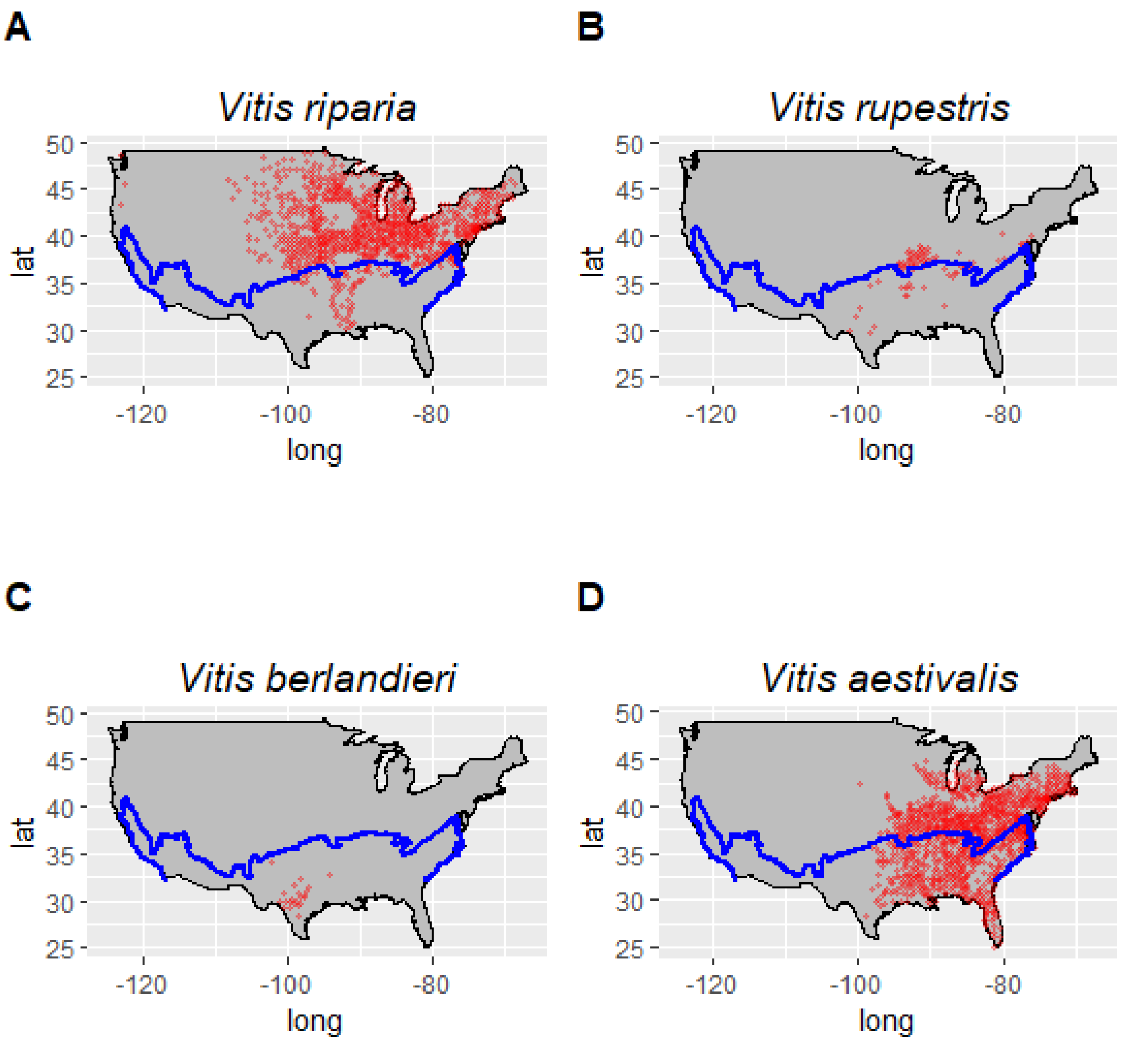


**Figure S11.** Distribution maps of the four main native American *Vitis* species. Blue line represents the risk index, $r > 0$, boundary for Pierce's disease. Red crosses recorded locations.

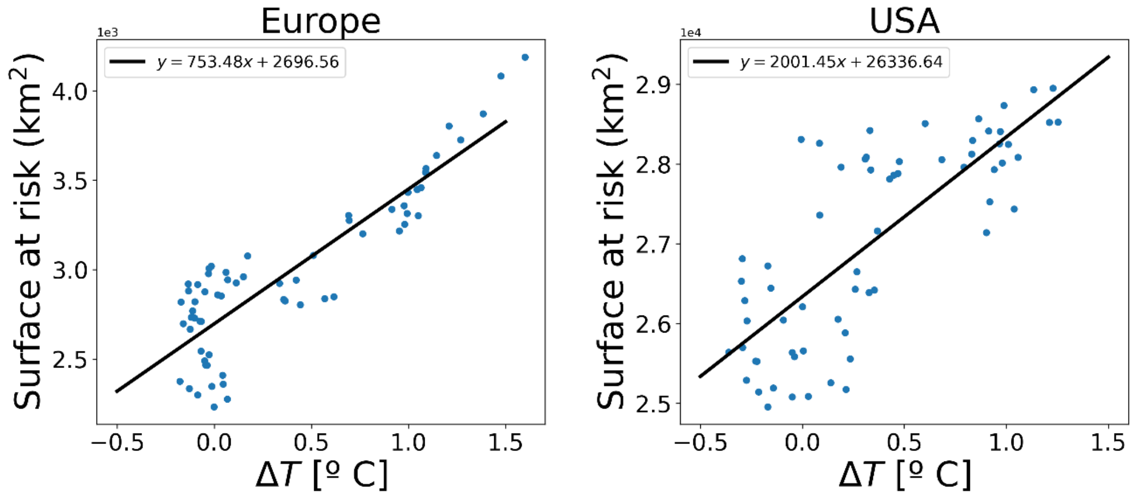


**Figure S12**. Relationship between temperature anomalies respect to the average (1850-1890 period) and the extension of PD in Europe (a) and the USA. (left) Risk areas in continental Europe contracts abruptly as there is no area in risk below 0 anomaly in comparison to the USA where the contraction is progressive (right).

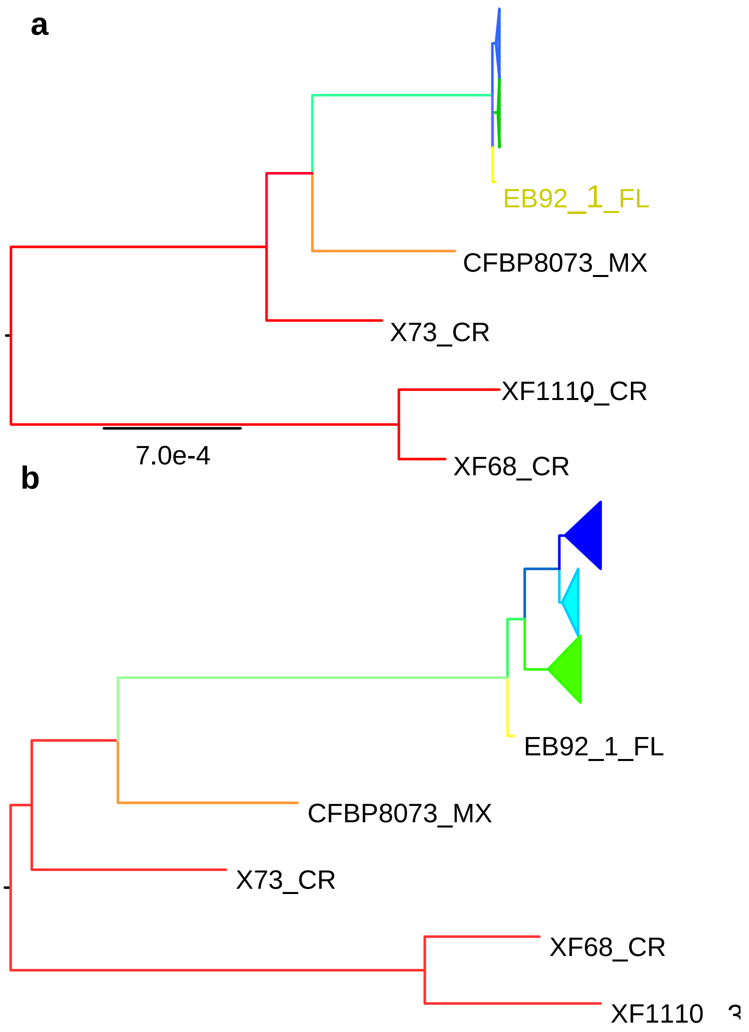


**Figure S13.** The effect of recombination in tree topology. (**a**) RAxML recombination-free tree showing the genetic position of the WUCC and EUCC respect *Xylella fastidiosa* subsp. *fastidiosa* isolates from Central America (red line). California clade (blue) and the EUCC (green) are collapsed. EB92-1 from Florida shows recombination with the subspecies *multiplex* and therefore was excluded from the $Xf_{PD}$ lineage. (b) *Xylella fastidiosa* subsp. *fastidiosa* tree before removing recombinant sequences.

**Supplementary Tables**

**Table S1.** Marginal likelihood estimation (MLE) of twelve models in the Bayesian phylogenetic analysis for the Southeastern US, Taiwan and the global phylogenetic tree. The five best-fitting models are marked in grey.

***Southeastern US***

| Substitution Model | clock model | Demographic model | Marginal |
|---|---|---|---|
| HKY | Strict | Bayesian Skyline | -2103570.90 |
| HKY | Strict | Coalescent constant | -2103575.12 |
| GTR | Strict | Coalescent constant | -2103575.32 |
| HKY | Strict | Exponential coalescent | -2103576.44 |
| HKY | UCLN | Bayesian Skyline | -2103583.81 |
| GTR | UCLN | Coalescent constant | -2103587.88 |
| GTR | Strict | Bayesian Skyline | -2103603.11 |
| GTR | UCLN | Bayesian Skyline | -2103607.05 |
| GTR | Strict | Exponential coalescent | -2103607.64 |
| HKY | UCLN | Coalescent constant | -2103607.81 |
| HKY | UCLN | Exponential coalescent | -2103608.79 |
| GTR | UCLN | Exponential coalescent | -2103611.86 |

***Taiwan***

| Substitution Model | clock model | Demographic model | Marginal |
|---|---|---|---|
| GTR | Strict | Bayesian Skyline | -2103385.58 |
| HKY | Strict | Exponential coalescent | -2103386.81 |
| GTR | UCLN | Coalescent constant | -2103390.20 |
| HKY | Strict | Coalescent constant | -2103390.80 |
| HKY | Strict | Bayesian Skyline | -2103391.73 |
| HKY | UCLN | Coalescent constant | -2103393.08 |
| GTR | Strict | Coalescent constant | -2103393.54 |
| GTR | UCLN | Bayesian Skyline | -2103451.13 |
| HKY | UCLN | Bayesian Skyline | -2103451.54 |
| GTR | UCLN | Exponential coalescent | -2103455.24 |
| HKY | UCLN | Exponential coalescent | -2103457.71 |
| GTR | Strict | Exponential coalescent | -2103465.46 |

***Whole tree***

| Substitution Model | clock model | Demographic model | Marginal |
|---|---|---|---|
| GTR | UCLN | Coalescent constant | -2113167.44 |
| GTR | UCLN | Exponential coalescent | -2113167.95 |
| HKY | UCLN | Coalescent constant | -2113191.08 |

| | | | |
|---|---|---|---|
| HKY | UCLN | Exponential coalescent | -2113193.93 |
| HKY | UCLN | Bayesian Skyline | -2113204.86 |
| GTR | Strict | Bayesian Skyline | -2113219.13 |
| GTR | Strict | Exponential coalescent | -2113222.57 |
| GTR | Strict | Coalescent constant | -2113224.62 |
| HKY | Strict | Exponential coalescent | -2113241.75 |
| HKY | Strict | Coalescent constant | -2113248.51 |
| HKY | Strict | Bayesian Skyline | -2113247.26 |
| GTR | UCLN | Bayesian Skyline | -2113697.90 |

**Table S2.** Summary of results in the inoculation test with American vines and hybrids.

| Rootstock | Origin | Nº leaves | DI | AUDCP | % Positive |
|---|---|---|---|---|---|
| 196-17 | (*V. vinifera* x *V. rupestris*) *x V. riparia* | 11.6 ± 4.2 | 4.33 | 22.89 | 100 |
| P1103 | *V. berlandieri* x *V. rupestris* | 5.7 ± 4.4 | 2.67 | 10.33 | 72.22 |
| R140 | *V. V. berlandieri* x *V. rupestris* | 5.3 ± 4.2 | 2.51 | 8.55 | 62.78 |
| 41B | *V. vinifera* X *V. berlandieri* | 3.6 ± 2.4 | 1.89 | 8.89 | 61.11 |
| 196-17(E23) | (*V. vinifera* x *V. rupertris*) x *V. riparia* | 3.9 ± 5.5 | 1.78 | 6.34 | 33.33 |
| R140 | *V. berlandieri* x *V. rupestris* | 2.7 ± 2.0 | 1.56 | 6.67 | 44.44 |
| R110 | *V. berlandieri* x *V. rupestris* | 2.5 ± 8.4 | 1.50 | 4.39 | 83.33 |
| 41B | *V. vinifera* X *V. berlandieri* | 2.2 ± 3.6 | 1.11 | 3.33 | 44.44 |
| P1103 | *V. berlandieri* x *V. rupestris* | 1.7 ± 2.0 | 1.11 | 4.34 | 0.00 |
| R110 | *V. berlandieri* x *V. rupestris* | 1.0 ± 1.0 | 0.78 | 3.88 | 26.00 |

Legends for Movies S1 to S5

**Movie 1**. PD risk in Europe and North America between 1850 and 1992 using coarse climate data. Maps are based on homogenous distribution of the vector and a $R_0$=5.

**Movie 2**. PD risk in Europe between 1950 to 2020 using ERA5-Land hourly climatic data and taking into account the distribution of the main vector *Philaenus spumarius*.

**Movie 3**. Annual risk maps from 1850 to 1992 based on model calibration with data obtained from inoculation trials on American *Vitis* species and hybrids. The model is run assuming a homogeneous distribution of vectors and a $R_0$=5.

**Movie 4.** Evolution of PD risk in Mediterranean southern France from 1950 to 2024.

**Movie 5**. Evolution of PD risk in the Iberian Peninsula (Spain and Portugal) from 1980 to 2021 using high resolution climatic data from Chelsa.

Legends for Datasets

**Dataset S1.** List of the genomes of *Xylella fastidiosa* isolates used in the study.

**Dataset S2**. Average Nucleotide Identity (ANI) based on complete assembled genomes of *Xylella fastidiosa* subsp*. fastidiosa* (2.4Mb) available in NCBI as of February 2004 using Jspecies.
